# Meiotic DSB-independent role of protein phosphatase 4 in Hop1 assembly to promote meiotic chromosome axis formation in budding yeast

**DOI:** 10.1101/2021.05.10.443451

**Authors:** Ke Li, Miki Shinohara

## Abstract

Dynamic changes in chromosomal structure that occur during meiotic prophase play an important role in the progression of meiosis. Among these structures, meiosis-specific chromosomal axis-loop structures are important as a scaffold for integrated control between the meiotic recombination reaction and the associated checkpoint system to ensure accurate chromosome segregation. However, the molecular mechanism of the initial step of chromosome axis-loop construction is not well understood. Here, we showed that in budding yeast, protein phosphatase 4 (PP4) that primarily counteracts Mec1/Tel1 phosphorylation is required to promote the assembly of chromosomal axis components Hop1 and Red1 onto meiotic chromatin via interaction with Hop1. PP4, in contrast, did not affect Rec8 assembly. Notably, unlike the previously known function of PP4, this novel function of PP4 was independent of meiotic DSB-dependent Tel1/Mec1 kinase activities. The defect in Hop1/Red1 assembly in the absence of PP4 function was not suppressed by dysfunction of Pch2, which removes Hop1 protein from the chromosome axis. This suggests that PP4 is required for the initial step of chromatin loading of Hop1 rather than stabilization of Hop1 on axes. These results indicate that phosphorylation/dephosphorylation-mediated regulation of Hop1 recruitment onto chromatin occurs during chromosome axis construction before meiotic double-strand break formation.

## Introduction

Meiotic recombination is essential for proper segregation of homologous chromosomes during the first meiotic division (meiosis-I). This process is initiated by Spo11-dependent programmed meiotic DNA double-strand break (DSB) formation at meiotic recombination hot spots (Keeney 2001). Meiotic DSB formation and subsequent meiotic recombination require the specific scaffold of an axis-loop structure of chromatin (Shinohara *et al*. 2000; Blat *et al*. 2002; Panizza *et al*. 2011).The highly conserved meiotic chromosome axis structure contains core factors, Red1, Hop1, and meiosis-specific cohesin complexes, including the meiosis-specific subunit Rec8 in budding yeast (Rockmill And Roeder 1988; Hollingsworth And Byers 1989; Klein *et al*. 1999). After meiotic DSB formation, recombinases Rad51/Dmc1 are recruited to single-stranded DSB ends (Bishop *et al*. 1992; Bishop 1994; Shinohara *et al*. 1997). The single-stranded DSB ends promote the assembly of Zip-Msh-Mer/synapsis initiation complex (ZMM/SIC) proteins, including the transverse element component Zip1, to construct the synaptonemal complex (SC) and promote crossover (CO) formation (Sym *et al*. 1993; Novak *et al*. 2001; Borner *et al*. 2004; Shinohara *et al*. 2008; Shinohara *et al*. 2015; De Muyt *et al*. 2018). In the early leptotene, Zip1 with Zip3 localizes to the centromere and mediates centromere pairing, an initial process that ultimately promotes chromosomal pairing (Tsubouchi and Roeder 2005; Falk *et al*. 2010).

Hop1 is an axis-associated HORMA-domain protein that plays multiple roles during meiotic recombination. Hop1 is required for efficient meiotic DSB formation, inter-homolog bias in meiotic recombination, and meiotic prophase checkpoint activation and inactivation (Hollingsworth *et al*. 1990; Schwacha and Kleckner 1997; Niu *et al*. 2005; Carballo *et al*. 2008; Panizza *et al*. 2011; West *et al*. 2018).

Hop1 together with Red1 promotes DSB formation. Hop1 is phosphorylated by Tel1/Mec1 (yeast ATM/ATR) kinases. These DSB sensor kinases activate the meiotic prophase checkpoint signaling pathway via Mek1/Mre4 kinase in response to meiotic recombination and synapsis defects (Niu *et al*. 2005; Carballo *et al*. 2008; Ho and Burgess 2011; Penedos *et al*. 2015; Herruzo *et al*. 2016). Hop1, Red1, and Mek1/Mre4 form the RHM complex involved in forming a stable axis structural scaffold for integrating recombination, as well as the required regulatory responses associated therewith. During meiotic DSB repair, homolog synapsis, mediated by SC formation, and a conserved AAA+ ATPase Pch2 promote Hop1 removal from the synapsed chromosome axis and attenuate the checkpoint signal (Borner *et al*. 2008; Herruzo *et al*. 2016). Recently, Pch2 was demonstrated to be involved in the activation, as well as the inactivation, of meiotic prophase checkpoints through Hop1 (Raina and Vader 2020). Thus, Hop1 phosphorylation plays a key role in completing proper meiotic recombination.

Mec1 plays a major role in both maintaining genome integrity and meiotic recombination (Kato and Ogawa 1994; Weinert *et al*. 1994). Mec1 is an essential protein in budding yeast and *SML1* mutations can suppress the lethality of *mec1* disruption (Zhao *et al*. 1998). Tel1 and Mec1 (Tel1/Mec1) play such subtly redundant roles that the *tel1 mec1* double mutant lends to a senescent phenotype (Ritchie *et al*. 1999). Tel1/Mec1 is also required for integrated meiotic recombination and checkpoint activation. Mec1/Tel1 kinase intra-chromosomally regulates the distribution of meiotic DSB formation sites (Anderson *et al*. 2015; Joshi *et al*. 2015; Challa *et al*. 2019). It also regulates signal transduction in the meiotic DSB formative positive/negative feedback loop through Rec114 phosphorylation (Carballo *et al*. 2013). In addition, Mec1/Tel1 mediates both DSB interference as well as CO interference, thus lending to proper non-random distribution of meiotic COs (Zhang *et al*. 2011; Garcia *et al*. 2015; Shinohara *et al*. 2019).

Protein phosphatase 4 (PP4) is a conserved putative serine/threonine protein phosphatase. It functions specifically, in response to DNA damage, in attenuating the phosphorylation signal of Mec1 and Tel1 during mitosis and meiosis (Hoffmann *et al*. 1994; Keogh *et al*. 2006; O’neill *et al*. 2007; Falk *et al*. 2010; Lai *et al*. 2011; Hustedt *et al*. 2015). Mec1/Tel1 phosphorylates many target proteins including Rad53, an ortholog of Chk2 kinase in budding yeast. In the mitotic DNA damage response pathway, PP4 dephosphorylates these targets in the process of recovery from DNA damage-induced signal transduction (O’neill *et al*. 2007). Several studies have revealed the meiotic DSB-associated function of PP4 during meiotic recombination (Falk *et al*. 2010; Chuang *et al*. 2012). During meiosis, centromere pairing is regulated by the balance of phosphorylation and dephosphorylation of Zip1 through Mec1/Tel1 kinase and PP4 phosphatase (Falk *et al*. 2010).

Here, we focused on the novel function of PP4, which works independently of meiotic DSB formation and its associated Mec1/Tel1 activity in early prophase-I. We were especially interested in the mechanics of its proper meiotic chromosome axis construction via physical interaction with Hop1 protein.

## Results

### PP4 activity is required for timely meiotic DSB formation and efficient CO formation

Pph3 is the catalytic subunit of the yeast PP4 complex. Pph3 contains several serine/threonine phosphatase motifs (Hoffmann *et al*. 1994). We constructed a catalytic mutation in the yeast *PPH3* gene by substituting the conserved histidine residue 112 required for phosphate binding with asparagine (Shi 2009), to generate the *pph3-H112N* mutant (Figure 1A). It is reported that Pph3 is required for Rad53 dephosphorylation and DNA damage repair in mitotic cells (O’neill *et al*. 2007). The *pph3-H112N* mutation did not affect the protein production of another PP4 subunit Psy2, which was analyzed with conjugation of a 13-repeated Myc epitope tag (Psy2-13Myc) (Figure S1A). In addition, we analyzed the assembly of Psy2-13Myc on meiotic chromatin in pachytene cells (Figure 1B). Consistent with a previous study (Falk *et al*. 2010), we detected Psy2-13Myc signals as punctate foci on meiotic chromatin.

**Figure1.**
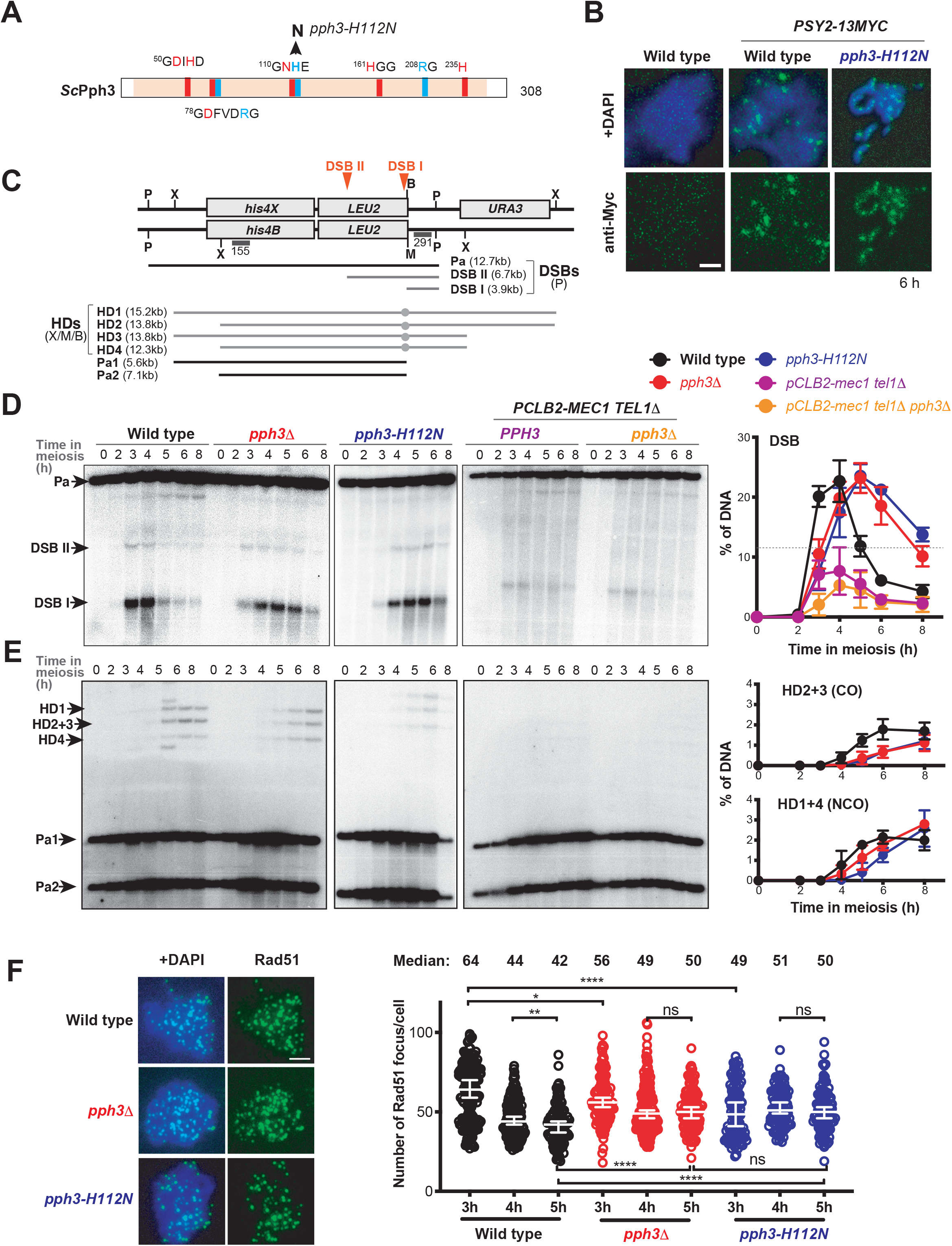
PP4 dephosphorylation activity is required for proper meiotic recombination. (A) Schematic of the functional domain structure of yeast Pph3 (*sc*Pph3) protein. Amino acid sequences contributing to phosphate binding and metal coordination are shown in pale blue and red, respectively. The position of the *pph3-H112N* and the substitution of histidine 112 with asparagine is shown. (B) Localization of Psy2-13Myc was assessed in *PPH3* (MSY6198/6190) and the *pph3-H112N* (MSY6242/6244) background was assessed via immunostaining with anti-Myc antibody (green). Representative images at 6h in meiosis are shown. Scale bar indicates 2 µm. (C) Schematics of the *HIS4-LEU2* recombination hotspot. Diagnostic P: *Pst*I, X: *Xho*I, B: *Bam*HI and M: *Mlu*I restriction sites are shown. The size of meiotic DSBs (DSB I, DSB II), and parental (Pa), and heteroduplexes (HDs) that are associated with COs (HD2 and HD3) or NCOs (HD1 and HD4), Parental (Pa1 and Pa2), and the positions of detectable probes 291 and 155 are shown. (D) Representative Southern blotting of DSB detection in wild type (black, NKY1303/1543), *pph3*Δ (red, MSY5632/MSY5634), *pph3-H112N* (blue, MSY6219/6220), p*CLB2*-*MEC1 tel1*Δ (purple, MSY5544/5546), and p*CLB2*-*MEC1 tel1*Δ *pph3*Δ (orange, MSY5760/5762) using probe 291. The graph shows the mean and standard error of the quantified DSB I signals from more than three reperated trials. (E) Representative Southern blotting of HD-associated COs and NCOs formed at the *HIS4-LEU2* hotspot using probe155. The upper graph shows the mean and standard error of the quantified HD-associated COs (HD2 and HD3). The lower graph shows the mean and standard error of the quantified HD-associated NCOs (HD1 and HD4) of wild type, *pph3*Δ, and *pph3-H112N*. (F) Representative immunostaining images of Rad51 (green) at 3 h on meiotic nuclear spreads in wild type (NKY1303/1543), *pph3*Δ (MSY5632/MSY5634), and *pph3-H112N* (MSY6219/6220). Scale bar indicates 2 µm. The dot plot graph shows the distribution of the number of Rad51 foci in each Rad51 focus positive nucleus at the indicated time points in wild type (black, NKY1303/1543), *pph3*Δ (red, MSY5632/MSY5634), and *pph3-H112N* (blue, MSY6219/6220). The data are presented as the medians and 95% confidence interval. Asterisks indicate statistically significant differences between the indicated strains determined using the Mann-Whitney U-test (* *P*<0.05, ** *P*<0.01, **** *P*<0.0001) using Prism9 software.

Importantly, the *pph3-H112N* mutation did not affect the Psy2-13Myc localization. Pph3 and Psy2 form a very stable PP4 complex (O’neill *et al*. 2007), thus suggesting that the catalytic activity of PP4 is not required for its meiotic chromatin localization. Falk et al. already reported the meiotic phenotypes of the *pph3*-null (*pph3*Δ) mutant (Falk *et al*. 2010). Therefore, we characterized two *pph3* mutants, *pph3*Δ and *pph3-H112N*, to better understand the function of PP4 and its phosphatase activity during meiosis. First, it was revealed that both *pph3* mutants showed similarly delayed meiosis progression compared to the wild type (Figure S1B). This finding was consistent with a previous study on meiosis progression in the *pph3*Δ and *psy2*Δ mutants (Falk *et al*. 2010).

To confirm whether delayed meiosis progression in *pph3* mutants is caused by meiotic recombination defects, we analyzed programmed meiotic DSB formation as an initial event of meiotic recombination. Toward this end, Southern blotting was applied at the *HIS4-LEU2* hot spot in chromosome III in the *pph3* mutants, although meiotic crossover formation defects in *pph3*Δ have already been reported (Falk *et al*. 2010).

The *HIS4-LEU2* hot spot is a well-analyzed artificial meiotic DSB hotspot that includes two known sites, DSB I and II (Figure 1C). We detected and quantified DSB I and showed its frequency at each time point after meiotic entry (Figure 1D). Meiotic DSBs became detectable 3 h after meiosis entry, showed a peak at 4 h, and disappeared by 6 h in the wild type (Figure 1D). In contrast, both *pph3-H112N* and *pph3*Δ showed an approximate 1 h delay in DSB formation compared to that in the wild type at the time when the DSB reached half of the peak height (11%; Figure 1D, dashed line). In addition, the *pph3* mutants showed an accumulation of meiotic DSBs with hyper-resected ends until 6 h after meiosis entry, resulting in an approximately 3 h delay in DSB disappearance (Figure 1D). These observations indicated the necessity of PP4 phosphatase activity for timely DSB formation and turnover. It is reported that PP4-dependent Rad53 dephosphorylation stimulates DNA end resection in mitotic DNA damage response (Villoria *et al*. 2019). The opposite function of PP4 in the DSB end resection may be due to mitotic and meiotic differences in Rad53 activity (Cartagena-Lirola *et al*. 2008).

Previous studies have revealed that PP4 is required for meiotic centromere pairing. This was found to occur either through Zip1 dephosphorylation for Mec1-mediated phosphorylation at S75 (Keogh *et al*. 2006; Falk *et al*. 2010; Hustedt *et al*. 2015) or dephosphorylation of many other Mec1/Tel1 targets during DSB repair (Keogh *et al*. 2006; Hustedt *et al*. 2015). Thus, we constructed a mitotic cyclin *CLB2* promoter-driven *MEC1* (p*CLB2*-*MEC1*) *tel1*Δ allele to determine the effect of Mec1 and Tel1 on delayed DSB formation and/or turnover in the *pph3* mutants. It is known that p*CLB2* induces mitosis-specific gene expression. Thus, it can be inferred that p*CLB2*-*MEC1* causes meiosis-specific shutdown of *MEC1* transcription (Lee and Amon 2003). We confirmed that the effect of this construct is not different from that of the *mec1*Δ *sml1*Δ strain in meiotic Hop1 phosphorylation. In addition, p*CLB2*-*MEC1* showed significant lower spore viability (59.6 ± 4.6%) compared to the wild type (96.1 ± 1.8%, *P* < 0.0001). This was found to be indistinguishable from *mec1*Δ *sml1*Δ (55.5 ± 7.1%, *P* = 0.37) or other *MEC1*-defective alleles (Supplemental Figure 1A). The p*CLB2*-*MEC1 tel1*Δ cells showed a reduced level of meiotic DSB formation (7.7 ± 1.9% of total DNA) compared to that in the wild type (22.7 ± 3.1%). p*CLB2*-*MEC1 tel1*Δ *pph3*Δ triple mutant showed more severe reduction of DSB formation (5.3 ± 0.45%; Figure 1D, graph). At later time points, in contrast to the *pph3* mutants, the p*CLB2*-*MEC1 tel1*Δ and p*CLB2*-*MEC1 tel1*Δ *pph3*Δ mutants showed the timely disappearance of DSBs, i.e., 6 h into meiosis (Figure 1D). These findings suggested that DSB accumulation and the resulting delay in DSB repair, which was observed in the *pph3* mutants at later time points (at 5–6 h), was dependent on Mec1/Tel1 (Figure 1E). By contrast, in the initiation of DSB formation, the p*CLB2*-*MEC1 tel1*Δ *pph3*Δ triple mutant showed a peak at 4 h, whereas the p*CLB2-MEC1 tel1*Δ double mutant peaked between 3 and 4 h after onset of meiotic entry (Figure 1D, graph). As a delay in meiotic DSB formation in the absence of Tel1/Mec1 activity was observed, the delayed DSB formation caused by *pph3* mutations appeared to be independent of Mec1/Tel1.

Next, we analyzed hetero-duplex (HD) formation using Southern blot to examine meiotic inter-homolog recombination products using heteroallelic restriction enzyme sites *Mlu*I and *Bam*HI located near the DSB I site, and polymorphic *Xho*I sites around the *HIS4-LEU2* hot spot (Figure 1C) (Storlazzi *et al*. 1996). Corresponding to the delayed meiotic DSB appearance and disappearance at the hot spot, the formation of both HDs corresponding to intermediates for crossover (CO) and non-crossover (NCO) were delayed by 3 h in the *pph3* mutants relative to that in the wild type (Figure 1E). In addition, an increase of NCO intermediates (HD1 and 4) at 8 h and a slight reduction of CO intermediates (HD2 and 3) compared to that of the wild type were observed not only in the *pph3*Δ, as reported previously (Falk *et al*. 2010), but also in the *pph3-H112N* mutant (Figure 1E).

After DSB end resection, meiotic DSBs are recognized by recombinases Rad51 and Dmc1, which facilitate strand exchange within the formation of COs and NCOs. Cytological analysis revealed that Rad51appear transiently as punctate foci on meiotic chromatin structure in the wild type (Shinohara *et al*. 2000). Thus, we next examined the kinetics of Rad51 assembly and disassembly in the *pph3* mutants using cytological analysis (Figure 1F). Results indicated that, there was a 1 h delay in the Rad51 assembly among the *pph3* compared to that in the wild type. In addition, accumulation of Rad51 foci was evident at 5 h after meiotic entry (Supplemental Figure 1C). In contrast, a significantly lower number of Rad51 foci per nucleus was observed at an early time point (3 h) in the *pph3*Δ and *pph3-H112N* mutants compared to that in the wild type (64 ± 2.7, 56 ± 2.7, and 49 ± 3.3, respectively; Figure 1F). This further indicated that PP4, as well as its phosphatase activity, is required for the initiation of meiotic DSB formation in proper timing. In the disassembly phase of Rad51 foci, a significant reduction in the number of Rad51 foci per nucleus was observed in the wild type. However, continuous accumulation of Rad51 foci was observed in the *pph3* mutants corresponding to inefficient meiotic DSB repair.

### PP4 contributes to timely assembly of Hop1 onto chromatin independent from a meiotic DSB dependent Mec1/Tel1 activity

Hop1 is a meiosis-specific phosphorylation target of Mec1 and Tel1 (Carballo *et al*. 2008), and PP4 is involved in the dephosphorylation of Hop1 (Carballo *et al*. 2008; Falk *et al*. 2010). Furthermore, Hop1 is a HORMA-domain protein and contributes to the axis structure, in conjugation with Red1. Hop1 is essential for partner choice in meiotic recombination (known as inter-homolog bias), and for efficient DSB formation (Hollingsworth *et al*. 1990; Niu *et al*. 2005; Panizza *et al*. 2011; Subramanian *et al*. 2016). Since it was observed that PP4 is required for efficient initiation of meiotic DSB formation (Figure 1C and D), we analyzed the kinetics of Hop1 assembly onto chromatin during meiotic prophase-I (Figure 2A). In the wild type, Hop1 assembly began 2 h after the onset of meiosis, when is corresponding to leptotene and prior to DSB formation.

**Figure 2.**
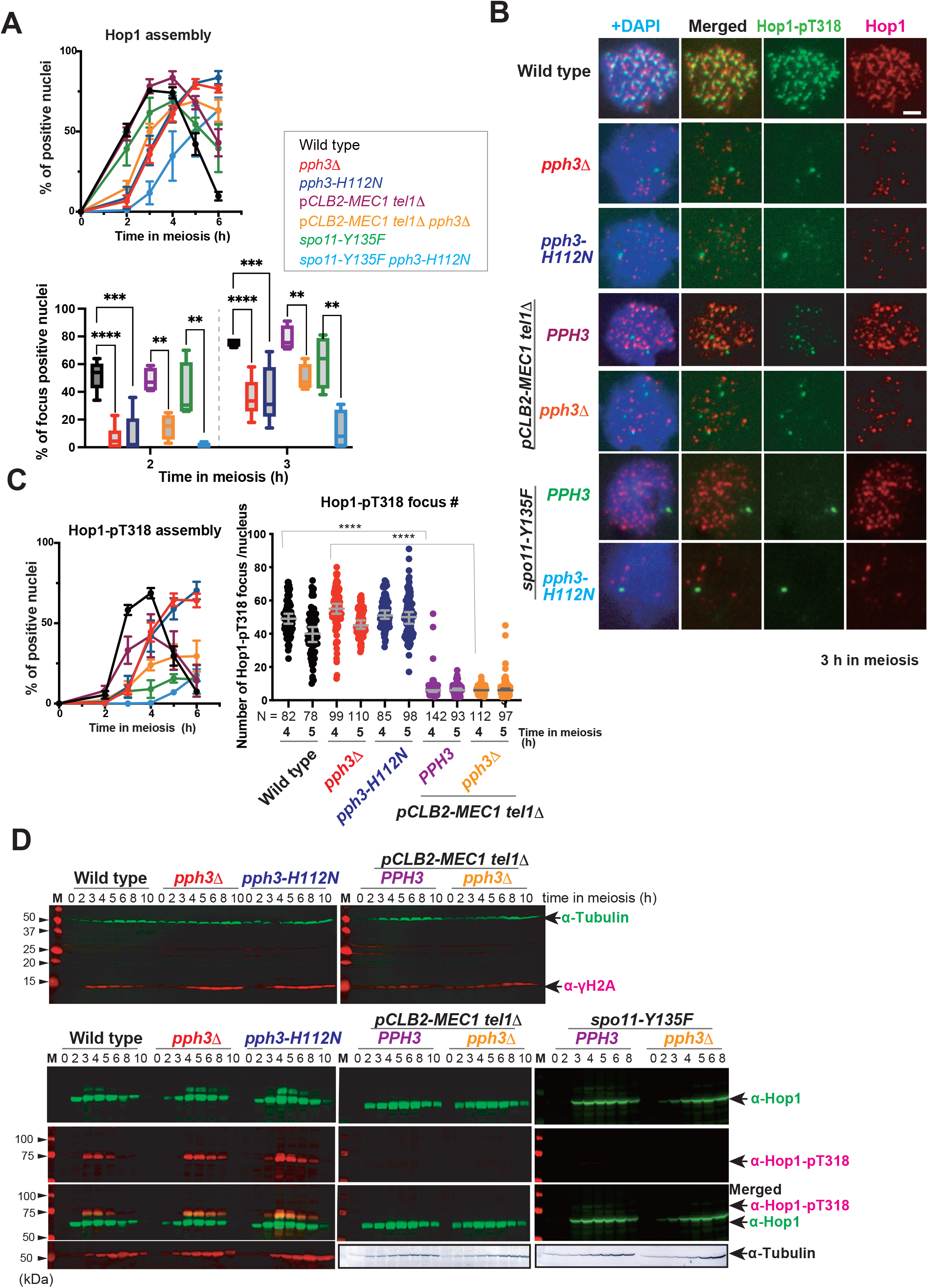
Hop1 assembly is delayed in the *pph3* mutant in early prophase-I. (A) Graphical depiction of the kinetics of Hop1 focus (upper) assembly on meiotic nuclear spreads in the indicated strains, wild type (black, NKY1303/1543), *pph3*Δ (red, MSY5632/MSY5634), *pph3-H112N* (blue, MSY6219/6220), p*CLB2*-*MEC1 tel1*Δ (purple, MSY5544/5546), p*CLB2*-*MEC1 tel1*Δ *pph3*Δ (orange, MSY5760/5762), *spo11-Y135F* (green, MSY4139/4141), and *spo11-Y135F pph3*-*H112N* (pale blue, MSY6568/6570). From the above graph, 2 h and 3 h data were extracted and re-presented as a box plot with statistical analysis. Whiskers show the median and interquartile range. More than 100 cells ere analyzed in each time point. Statistically significant differences were determined using Student’s t-test (\*\**P* < 0.001, *** *P* < 0.0005, **** *P* < 0.0001) using Prism9 software. (B) Representative cytological immunostaining images of Hop1 (green) and Hop1-pT318 (red) on meiotic nuclear spreads at 3 h in each strain shown in Figure 2A. Scale bar indicates 2 µm. (C) Graphical depiction of the kinetics of Hop1-pT318 focus (left) assembly on meiotic nuclear spreads in the indicated strains. The dot plot graph (right) shows the distributions of the number of Hop1-pT318 foci in each Hop1-pT318 focus positive nucleus at 4 and 5 h in wild type, *pph3*Δ, *pph3-H112N*, p*CLB2*-*MEC1 tel1*Δ, and p*CLB2*-*MEC1 tel1*Δ *pph3*Δ. The data are presented as the medians and 95% confidence interval. N denotes number analyzed. Statistically significant differences were determined using Mann-Whitney’s U-test (**** *P* < 0.0001) using Prism9 software. (D) Representative western blot images of phosphorylation of histone H2A (γH2A, red) and α-tubulin (green) at the indicated time point during meiosis (upper panel). Hop1 (green, top), phosphorylation of Hop1 at T318 (Hop1-pT318, red, middle), merged images of Hop1 and Hop1-pT318, and α-tubulin (bottom) at the indicated time points during meiosis. M denotes molecular weight markers.

We also analyzed the localization of Hop1, and Hop1 phosphorylation at threonine 318 (Hop1-pT318) on meiotic chromatin. Hop1-pT318 is dependent on Mec1/Tel1 kinase activation by Spo11-induced meiotic DSB formation (Carballo *et al*. 2008). Hop1-pT318 has been observed as punctate foci at the end of short patches of Hop1 signal in the wild type (Carballo *et al*. 2008) (Figure 2A and B). In the wild type, 50% of the cells displayed Hop1 foci 2 h after meiosis entry, but few Hop1-pT318 were observed at the same time. Hop1-pT318 foci began to appear after 3 h (Figure 2A and B). Although the formation of Hop1-pT318 foci was dependent upon Spo11 function, normal Hop1 assembly was also observed in the *spo11-Y135F*, which lacks the Spo11 catalytic tyrosine residue (Figure 2A and B). In addition, it is known that Hop1-pT318 focus formation requires single-stranded regions at meiotic DSB ends (Iwasaki *et al*. 2016). The approximately 2 h time lag evident between the Hop1 loading and the Hop1-pT318 signals observed in the present study indicate that, in wild type, Hop1 assembly occurs prior to the DSB formation and/or DSB-end resection. In *pph3* mutants, Hop1 assembly was rarely seen at 2 h compared to that in the wild type. After 2 h, Hop1 staining gradually increased, reaching the wild-type’s frequency (80%) at 5 h. Notably, pT318 staining appeared at approximately the same time (3 h) as the appearance of Hop1 foci (Figure 2A and B) and then gradually increased in ways similar to Hop1, suggesting that Hop1 loading seemed to be the rate-limiting factor for Hop1 phosphorylation in the *pph3* mutants. These results indicated that Pph3 and its activity can promote the timely loading of Hop1 on chromosomes. We further confirmed that this was not due to delayed expression of Hop1 in the mutants (see Figure 2C and Supplemental Figure 2B). In addition, the delayed assembly of Hop1 in the *pph3* mutants was observed in the *spo11-Y135F* mutation background (Figure 2A), thus indicating that PP4 promotes Hop1 assembly independent of meiotic DSB formation. Interestingly, Hop1 assembly in the *spo11-Y135F pph3-H112N* double mutant showed a significant delay compared to that in the *pph3-H112N* mutant (Figure 2A). This would suggest a parallel function of PP4 and Spo11 in Hop1 assembly or stabilization on the chromosome axes.

Spo11 independent PP4 function in Hop1 assembly suggests a possibility that meiotic DSB-dependent Mec1/Tel1 activation is not essential to PP4 function in Hop1 assembly. In contrast, the *pph3* mutant shows a Mec1-dependent meiotic DNA replication delay (Falk *et al*. 2010). Further, we confirmed delayed meiosis progression in the *spo11-Y135F pph3-H112N* double mutant as reported by Falk et al (Supplemental Figure 2A). We subsequently examined the effect of Mec1 and Tel1 on Hop1 assembly in *pph3*Δ mutants. Although p*CLB2*-*MEC1 tel1*Δ mutant cells did not display any defects in Hop1 assembly, the p*CLB2*-*MEC1 tel1*Δ *pph3*Δ mutant showed delayed Hop1 assembly, similar to the *pph3* mutants (Figure 2A and B). In contrast, although a few faint Hop1-pT318 foci were observed in the p*CLB2*-*MEC1 tel1*Δ mutation background, none were found in the *spo11-Y135F* mutation background (Figure 2B). This Hop1-pT318 focus formation raises doubts as to the presence of residual Mec1 activity in the p*CLB2*-*MEC1* meiosis-specific shout down allele. However, Hop1-pT318 foci in strains with p*CLB2*-*MEC1 tel1*Δ mutation backgrounds were qualitatively distinguishable from those in the strains with wild-type *MEC1* background (Figure 2C).

Next, we analyzed the phosphorylation status of target proteins of PP4 during meiosis using western blot. PP4 is known to be involved in dephosphorylation of S129 of histone H2A (γH2A) after DNA damage in vegetative growth cells (Keogh *et al*. 2006), as well as in Hop1-pT318 during meiosis (Chuang *et al*. 2012). Thus, we first confirmed that the γH2A signal accumulated in the *pph3* mutants during meiosis in a Mec1/Tel1 dependent manner, a pattern similar to that in mitotic cells, with markedly reduced γH2A signals in the p*CLB2*-*MEC1 tel1*Δ *pph3*Δ mutant (Figure 2D). Moreover, accumulation of Hop1-pT318 signal was observed until the 8 h point in the *pph3*Δ mutant. The presence of this phosphorylation depended upon Mec1/Tel1 (Figure 2D). Although Hop1 protein production was detectable from 2 h after meiosis entry in both wild-type and *pph3* mutants, it was slight delayed in the *pph3* mutants. By contrast, the initiation of Hop1 phosphorylation at T318 was delayed by 1 h in the *pph3* mutants before reaching maximum levels compared to that in the wild type. Notably, despite the timely early production of Hop1 protein in the *pph3* mutants (Figure 2D and Supplemental Figure 2B), the appearance of Hop1 foci on meiotic nuclear spread was significantly delayed in the *pph3* mutants (Figure 2A). This suggested that, although PP4 might not be largely involved in Hop1 production, it acted to promotes Hop1 loading onto chromatin. Moreover, Hop1 protein production kinetics were also normal in the p*CLB2*-*MEC1 tel1*Δ or p*CLB2*-*MEC1 tel1*Δ *pph3*Δ mutants. In contrast to finding for γH2A phosphorylation, residual Hop1-pT318 signal was not detected in the p*CLB2*-*MEC1 tel1*Δ background. This observation accorded with a previous report showing that Hop1 phosphorylation largely depends on Mec1 and Tel1 activities (Carballo *et al*. 2008). Overall, our findings indicate a defect in Hop1 assembly on chromatin, despite sufficient production of Hop1 protein.

### PP4 is involved in Red1-Hop1 complex assembly but not in meiotic cohesin assembly

Next, we examined Red1 and Rec8 meiotic chromosome axis components (Rockmill and Roeder 1988; Klein *et al*. 1999). Red1 forms a complex with Hop1 (Niu *et al*. 2005). Rec8 is a component of the meiosis-specific sister chromatid cohesin complex. Both proteins are required for the proper formation of both the meiotic axial elements, as well as a highly-ordered specific chromosome structure, termed the SC (Klein *et al*. 1999). Red1 and Rec8 are mutually independently recruited to chromatin (Sun *et al*. 2015). We examined Red1 and Rec8 staining on meiotic chromatin (Figure 3A). Wild type, Red1 and Rec8 colocalized with each other and formed linear structures along the 4′,6-diamidino-2-phenylindole (DAPI) nuclear signal on meiotic chromatin 3 h after meiosis entry. This co-stain pattern on Rec8 and Red1 differs slightly from previous reports (Kim *et al*. 2010). This may be due to differences in the antibodies used. By contrast, in the *pph3* mutants, Red1 and Rec8 were observed as many fine signals throughout the chromatin spreads. The number of Rec8 signals was clearly higher than that of Red1. Time course experiments revealed a significant delay in the assembly of Red1 protein in the *pph3* mutants, similar to the assembly of Hop1 shown in Figure 2A and 2B. However, the timing of Rec8 assembly, despite of its unique structure, was not affected in the presence or absence of Pph3 activity (Figure 3B). An additional delay in the disassembly of both Red1 and Rec8 was observed in the *pph3* mutants.

**Figure 3.**
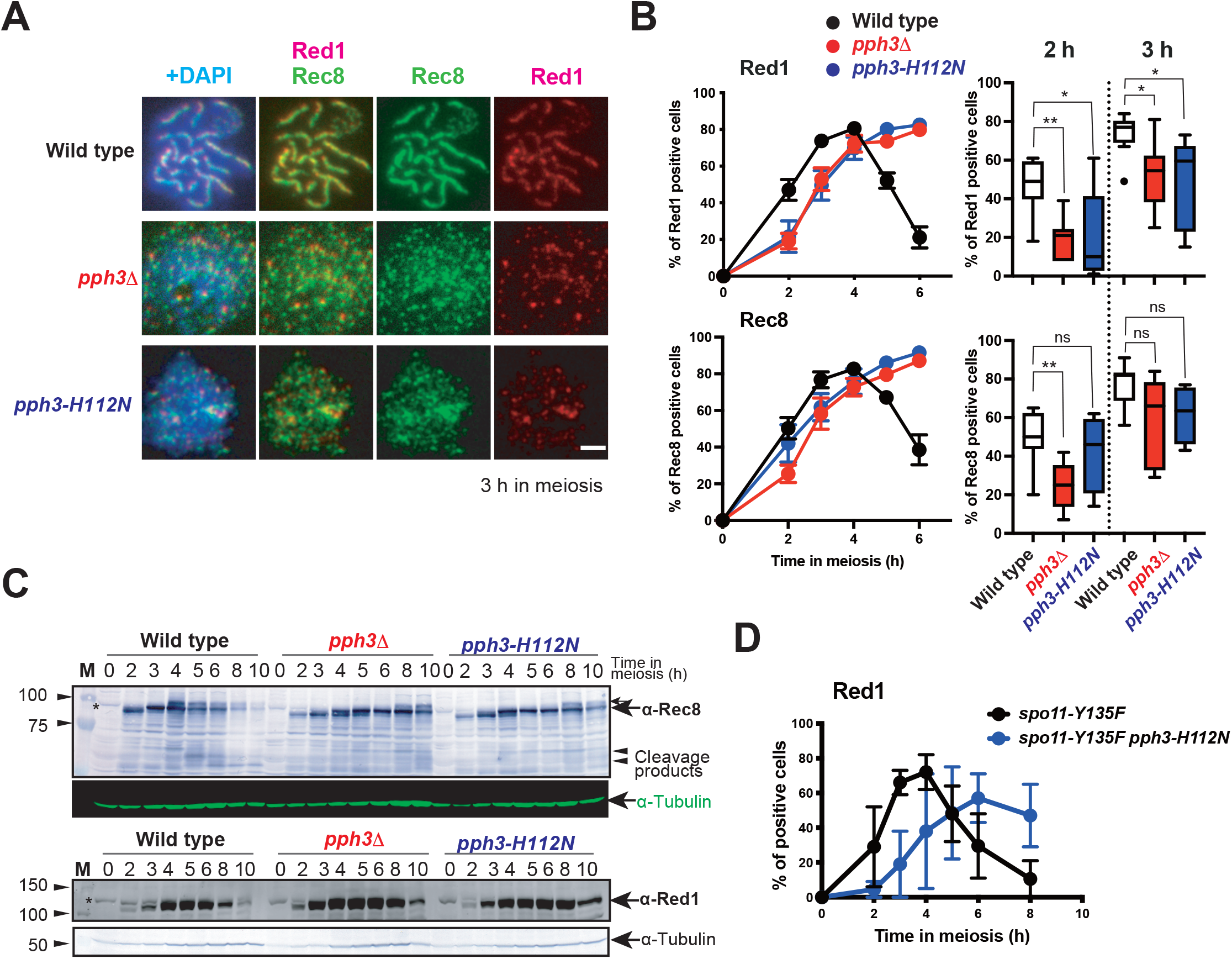
Rec8 appeared in normal timing in the *pph3* mutant in early prophase I. (A) Representative cytological immunostaining images of Rec8 (green) and Red1 (red) on meiotic nuclear spreads at 3 h in wild type (NKY1303/1543), *pph3*Δ (MSY5632/MSY5634), *pph3-H112N* (MSY6219/6220). wild type (NKY1303/1543), *pch2*Δ (MSY6446/6448), *pph3*Δ (MSY5632/MSY5634), and *pph3*Δ *pch2*Δ (MSY6417/6419) (B) Left graphs show the kinetics of Red1 (upper) and Rec8 (lower) focus assembly on meiotic nuclear spreads in the indicated strains, wild type (black, NKY1303/1543), *pph3*Δ (red, MSY5632/MSY5634), and *pph3-H112N* (blue, MSY6219/6220). More than 100 cells were analyzed at each time point. Error bars show the mean ± standard deviation of more than three trial repetitions. Right box whiskers plots show distributions of frequencies (%) of nuclei positive for Red1 (upper) or Rec8 (lower) at 2 or 3 h from the data sets shown in the left graphs. Whiskers show the median and interquartile range. Statistically significant differences were determined using Welch’s t-test (* *P* < 0.05, ** P < 0.01) using Prism9 software. (C) Representative western blot images of Rec8 (upper) and Red1 (lower), with α-tubulin images as internal controls, at the indicated meiotic time points. M denotes molecular weight markers. Asterisks indicate non-specific protein signals. (D) Graphical depiction of the kinetics of Red1 focus assembly on meiotic nuclear spreads in the *spo11-Y135F* (black, MSY4139/4141) and *spo11-Y135F pph3*-*H112N* (blue, MSY6568/6570). More than 100 cells were analyzed in each time point. Error bars show the mean ± standard deviation of more than three repeated trials.

Next, we analyzed Red1 and Rec8 protein production during meiosis using western blot (Figure 3C). Rec8 phosphorylation via Dbf4-dependent kinase (DDK) and polo-like kinase 1 (PLK1) is associated with cleavage-independent cohesion release from late prophase-I chromatin (Challa *et al*. 2019). Although Rec8 appeared at approximately the same time in the *pph3* mutants as in the wild type, the phosphorylation of Rec8 was delayed in the *pph3* mutants compared to in the wild type. Corresponding to delayed Rec8 phosphorylation, Rec8 cleavage that occurs during the meiotic divisions was also delayed in the *pph3* mutants. These results were consistent with delayed meiosis progression in *pph3* mutants (Supplemental Figure 2A). In addition, normal levels of Red1 protein were observed in *pph3* mutants (Figure 3C). These results suggest that PP4 is essential to properly timed Hop1-Red1 loading, but not Rec8 cohesin, which is solely required for basic axis structure construction.

### PP4 physically interacts with Hop1 protein

We next examined physical interactions between the Hop1-Red1 and PP4 using a co-immunoprecipitation assay (Figure 4A). Psy2-13Myc protein was precipitated using anti-Myc antibody in meiotic cell extracts of *PSY2-13MYC* derivative *PPH3* (wild type) cells. The Hop1 signal was detected in the whole cell extracts (WCEs) at 2 to 6 h during meiosis. After immunoprecipitation in *PSY2-13MYC* cell extracts, the Hop1 protein signal was observed at 2 and 4 h after meiosis entry, indicating the interaction of Pys2 with Hop1. However, despite of the presence of Hop1 protein in the WCEs, the signal was not apparent at 6 h. Since Hop1-PP4 physical interaction was detected, we analyzed other axis component Red1 and Rec8 proteins in the precipitates using western blot. By contrast, Red1 and Rec8 were not observed in the precipitates, even though both proteins were present in the WCEs. The findings indicate that PP4 primarily interacts with Hop1, but not Red1 or Rec8. Given that Hop1 binds to Red1 as a complex (Niu *et al*. 2005), there might be two distinct populations of Hop1, Hop1-PP4 and Hop1-Red1.

**Figure 4.**
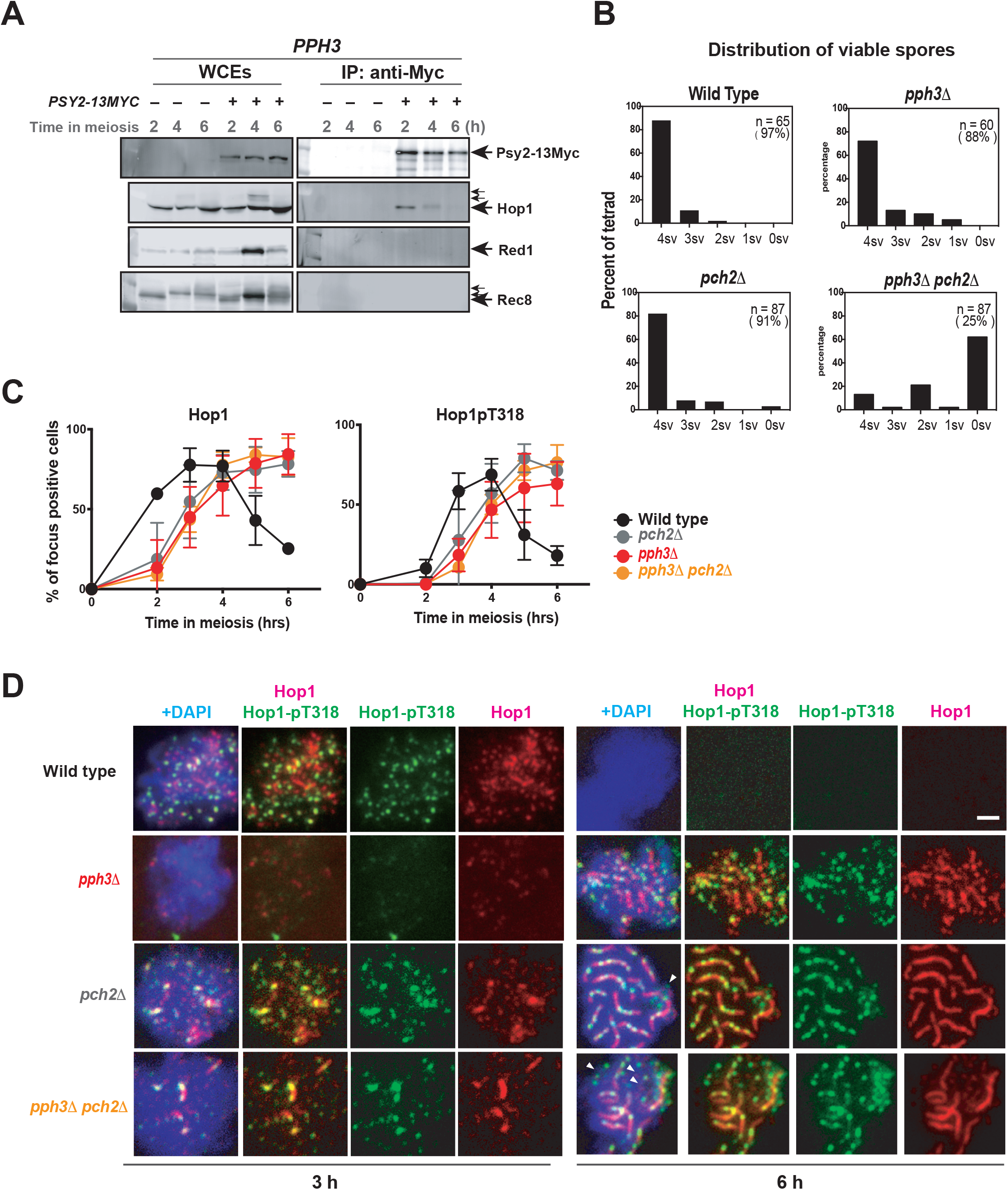
PP4 interacts with Hop1 during meiosis. (A) Immunoprecipitation (IP) analysis of Psy2-13Myc from *PPH3* (non-tag, NKY1303/1543) and *PPH3* (*PSY2-13MYC*, MSY6188/6190) cells at the indicated time points during meiosis. Whole cell extracts (WCEs) and IP products were probed with anti-Myc, anti-Hop1, anti-Red1 and anti-Rec8 antibodies. Small arrows show the modified products. (B) Tetrad analysis of spore viability in wild type (NKY1303/1543), *pph3*Δ (MSY5632/MSY5634), *pch2*Δ (MSY6446/6448), and *pph3*Δ *pch2*Δ (MSY6417/6419). The graphs show the fraction of tetrads containing 0-, 1-, 2-, 3- and 4-viable spores. The overall spore viability is shown in parentheses. n: number of asci analyzed. (C) Graphical depiction of the kinetics of Hop1 (left) and Hop1-pT318 (right) focus assembly on meiotic nuclear spreads in the indicated strains: wild type (black, NKY1303/1543), *pch2*Δ (gray, MSY6446/6448), *pph3*Δ (red, MSY5632/MSY5634), and *pph3*Δ *pch2*Δ (orange, MSY6417/6419). More than 100 cells were analyzed per measured time point. Error bars show the mean ± standard deviation of more than three repeated trials. (D) Representative cytological immunostaining images of Hop1 (green) and Hop1-pT318 (red) on meiotic nuclear spreads at 3 h (left panel) and 6 h (right panel) in wild type (NKY1303/1543), *pch2*Δ (MSY6446/6448), *pph3*Δ (MSY5632/MSY5634), and *pph3*Δ *pch2*Δ (MSY6417/6419). Arrowheads show unsynapsed regions. Scale bar indicates 2 µm.

### Defective Hop1 assembly onto chromosome axes in the *pch2*Δ background in the *pph3* mutant

Pch2 is a AAA+ ATPase that promotes Hop1 turnover from synapsed chromosome axes. It is known to be abundant localization of Hop1 protein on the synapsed axes in *pch2*Δ cells. Pch2 is also involved in a Hop1 dephosphorylation (Lo *et al*. 2014; Herruzo *et al*. 2016; Subramanian *et al*. 2016). Given our observation a defect in Hop1 assembly and delayed phosphorylation of Hop1 in the *pph3* mutants, we analyzed the effects of the *pph3* mutations in the Hop1 assembly onto meiotic chromosome axis in the *pch2*Δ background.

First, spore viability and distribution of viable spores were analyzed in the *pph3*Δ *pch2*Δ double mutant using tetrad analysis (Figure 4B). Spore viability in the wild type, *pph3*Δ, and *pch2*Δ was 97%, 88%, and 91%, respectively. Both *pph3*Δ and *pch2*Δ had slightly reduced spore viability, but there was no significant bias in the distribution of viable spores in asci, as reported previously (San-Segundo and Roeder 1999; Falk *et al*. 2010). By contrast, the spore viability of the *pph3*Δ *pch2*Δ double mutant was only 25%, with distributions of 0-, 2-, and 4-viable spores. This bias was indicative of the non-disjunction of homologous chromosomes in the first meiotic division.

In Hop1 and Hop1-pT318 assembly/disassembly kinetics, Hop1-pT318 assembly was delayed in the *pch2*Δ mutant, consistent with a delayed meiotic DSB formation (Farmer *et al*. 2012). This delay was similar to that in the *pph3*Δ and the *pph3*Δ *pch2*Δ double mutants (Figure 4C and D). Thus, indicating the absence of additive defects to *pph3*Δ in Hop1 assembly/disassembly in the *pch2*Δ mutation. The findings suggest that Pch2 and PP4 function in the same pathway in Hop1 assembly/disassembly.

Next, we examined the effect of the *pch2*Δ mutation on Hop1 localization by using cytological analysis. As indicated previously (IWASAKI *et al*. 2016) and in Figure 2A, Hop1-pT318 foci were observed as punctate foci in the wild type and were affected in the *pph3*Δ in early prophase-I (approximately 3 h in meiosis) (Figure 4D). In the *pch2*Δ mutant, Hop1 aggregates colocalized with Hop1-pT318 signals. Similar aggregates were noted in the *pph3*Δ *pch2*Δ double mutant at an early time point.

Notably, at the later time point (6 h during meiosis), Hop1-pT318 signals were observed as a beaded chain on the fully elongated Hop1 signal on meiotic chromosomes in the *pch2*Δ mutant cells (Figure 4D). In the *pph3*Δ *pch2*Δ double mutant, some Hop1 signals were observed as dots and some were colocalized with the Hop1-pT318 signal, similarly to that in *pph3*Δ. These findings suggest that the *pph3*Δ mutation is epistatic to the *pch2*Δ mutation in Hop1 localization process. This further indicates that PP4 may function to promote Hop1 assembly onto meiotic chromatin that lends to construction of the chromosome axis, a process occurring prior to Pch2 function in Hop1 removal from the chromosome axis. By contrast, in the *pph3*Δ *pch2*Δ double mutant, some Hop1-pT318 signals were observed as beaded chain on fully elongated Hop1. This indicates that the *pch2*Δ mutation stabilized Hop1 protein in the *pph3*Δ on some chromosome axis, but not on others. Thus, other pathways may be able to promote Hop1 assembly in the absence of PP4.

### Multiple pathways to promote Hop1 assembly

On the meiotic chromosome axes, Hop1 is dynamic and is removed from chromosomes through Pch2 activity. Then, Hop1 is ultimately replaced by the transverse element component Zip1 in the pachytene (Borner *et al*. 2008). Given this, we examined Zip1 localization in the *pph3*Δ *pch2*Δ double mutant. In wild-type cells, as previously reported, Zip1 was observed as dotty foci in the leptotene (2 h).It begins to elongate after the DSB end resection in the zygotene, followed by the extension of Zip1 throughout each chromosome in the pachytene (Sym *et al*. 1993; Storlazzi *et al*. 1996; Shinohara *et al*. 2015). Significant delays in both the appearance of dotty Zip1 signals and the Zip1 elongation were evident for *pph3*Δ (Figure 5A), as reported previously (Falk *et al*. 2010). The *pch2*Δ mutant showed Zip1 elongation delay, in addition to issociation from the chromosomes after full elongation. However, the elongated Zip1 became the major population at 5 h after meiosis entry. In the *pph3*Δ *pch2*Δ double mutant, after the delayed appearance of dotty Zip1 signals, similar to *pph3*Δ (2 to 3 h), compromised Zip1 elongation was observed. Complementary localization of Zip1 and Hop1 on chromosomes was observed from zygotene to pachytene in wild type. A similar localization pattern was evident in the *pph3*Δ mutant. By contrast, *pch2*Δ mutant cells showed uniform colocalization of Zip1 and Hop1 (Figure 5B), as previously reported (BORNER *et al*. 2008). *Saccharomyces cerevisiae* haploid cells have 16 chromosomes. Theoretically, 16 bivalents should be observed in the pachytene stage. However, the number of linear Zip1 signals per cell was 12 ± 2.7 (mean ± 95% confidence interval) in the *pch2*Δ. This amout was significantly reduced in the *pph3*Δ *pch2*Δ double mutant in 68% of *pch2*Δ (8.2 ± 2.6) (Figure 5B and C).

**Figure 5.**
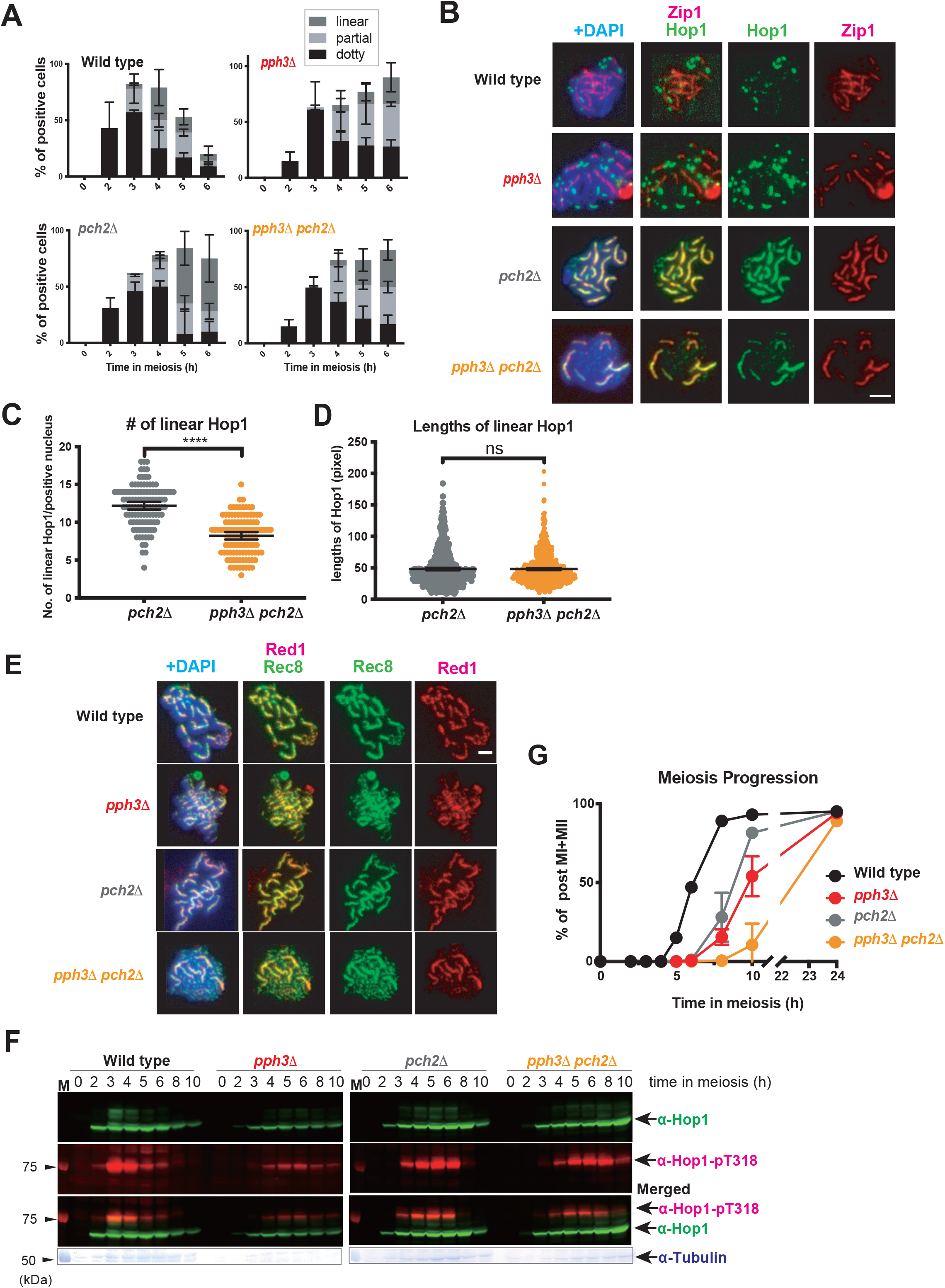
*pch2*Δ does not completely suppress the defect in Hop1 assembly of *pph3*Δ. (A) Zip1 and Hop1 elongation was analyzed in wild type (NKY1303/1543), *pch2*Δ (MSY6446/6448), *pph3*Δ (MSY5632/MSY5634), and *pph3*Δ *pch2*Δ (MSY6417/6419) via immunostaining at each time point. The graphs show the percentages of each class of Zip1 morphologies; linear (dark gray, pachytene), partial (pale gray, zygotene), and dotty (black, leptotene). Error bars show the mean ± standard deviation of more than three trials. More than 100 cells at each time point were analyzed. (B) Representative cytological immunostaining images of Hop1 (green) and Zip1 (red) on pachytene-phase nuclear spreads at 4 h in wild type (NKY1303/1543), and 6 h in the *pch2*Δ (MSY6446/6448), *pph3*Δ (MSY5632/MSY5634), and *pph3*Δ *pch2*Δ (MSY6417/6419). Scale bar indicates 2 µm. (C) The dot plot graph of the distribution of the number of linearized Hop1 signals in each pachytene stage nucleus in *pch2*Δ (N = 112) and *pph3*Δ *pch2*Δ (N = 109) at 6 h during meiosis from three independent trials. Error bars show the mean with 95% confidence interval. Statistically significant differences between the *pch2*Δ and *pph3*Δ *pch2*Δ were determined using the Mann-Whitney U-test (**** *P* < 0.0001) using Prism9 software. (D) The dot plot graph of the distribution of lengths of linearized Hop1 signal of pachytene stage nucleus in *pch2*Δ (gray, N = 718) and *pph3*Δ *pch2*Δ (orange, N = 780) at 6 h in meiosis from three independent trials. Error bars show the mean with 95% confidence interval. Statistically significant differences between the *pch2*Δ and *pph3*Δ *pch2*Δ were determined using the Mann-Whitney U-test (ns; not significant) using Prism9 software. (E) Representative cytological immunostaining images of Rec8 (green) and Red1 (red) on pachytene-stage nuclear spreads in wild type (3 h, NKY1303/1543), *pch2*Δ (6 h, MSY6446/6448), *pph3*Δ (6 h, MSY5632/MSY5634), and *pph3*Δ *pch2*Δ (6 h, MSY6417/6419). Scale bar indicates 2 µm. (F) Representative western blot images of Hop1 (green) as well as phosphorylation of Hop1 at pT318 (Hop1-pT318, red) and α-tubulin (bottom) at the indicated time points during meiosis. M denotes molecular weight markers. (G) Meiosis progression in wild type (black, NKY1303/1543), *pch2*Δ (gray, MSY6446/6448), *pph3*Δ (red, MSY5632/MSY5634), and *pph3*Δ *pch2*Δ (orange, MSY6417/6419) mutants are shown. Frequencies of nuclei with more than two DAPI-staining bodies at each time point are plotted. More than 100 cells were analyzed at each time point. Error bars show the means and standard deviation of two independent trials.

Although several elongated Zip1 signals colocalized with Hop1 in *pph3*Δ *pch2*Δ, Hop1-Zip1 free DAPI signals were also often observed in *pph3*Δ *pch2*Δ (Figure 5B). Next, we compared the length of a linear stretch of Zip1-Hop1 signals indicative of chromosome size in *pch2*Δ within *pph3*Δ *pch2*Δ, as specifically observed in the process of residual Zip1-Hop1 elongation in *pph3*Δ *pch2*Δ. No significant difference was evident (*P* = 0.19, Figure 5D).

The *pph3*Δ mutation showed a chromosome-dependent effect in Hop1-Zip1 elongation in the *pch2*Δ mutant background. We therefore analyzed Rec8 and Red1 localization, as alternative axis components, to determine the status of the Hop1-Zip1 free region observed in the *pph3*Δ *pch2*Δ double mutant. Unlike the Hop1 protein on the axes, Red1 is not therefrom by Pch2. In addition, Rec8 is recruited to chromatin independent of Hop1-Red1 (Sun *et al*. 2015). Further, Rec8 focus formation was not affected in the *pph3* mutants (Figure 3A). In *pch2*Δ mutant cell, Rec8 and Red1 evidently colocalized on the chromosome axes (Figure 5E). In contrast, in the *pph3*Δ *pch2*Δ double mutant, Rec8 and Red1 only colocalized on the synapsed chromosome axes. Further, many of the Rec8 was observed as a thin bead chain, where are not synapsed. These findings suggest that the *pph3* mutants are defective primarily at the Red1-Hop1 recruitment stage in the formation of the pre-Zip1 recruited chromosomal axis.

### Delayed meiosis progression in the *pph3*Δ occurs independent of Pch2

Hop1 is an adaptor of the meiosis-specific prophase checkpoint. Hop1 phosphorylation is affected in the *pch2*Δ mutant (Lo *et al*. 2014) and Mec1/Tel1-dependent phosphorylation of Hop1 affects Hop1 activity (Carballo *et al*. 2008). In the wild type, Hop1 phosphorylation was detected during meiosis starting at 3 h and continued until 5 h, upon which point, it disappeared (Figure 5F). The timing of this Hop1-pT318 disappearance aligned with the timing of meiosis-I initiation (Figure 5G).

This was also observed with *pph3*Δ mutant cells, in which the Hop1-pT318 signal disappeared at 8 h simultaneously with the appearance of post-meiotic cells. In *pch2*Δ mutant cells, consistent with previous findings, Hop1 protein accumulated until 10 h, and stimulated Hop1 phosphorylation disappeared by 8 h. In contrast, in the *pph3*Δ *pch2*Δ double mutant, Hop1 protein accumulated until 10 h. In turn, Hop1-pT318 signal also accumulated until 8 h after which it began to disappear. Notably, corresponding to the delayed disappearance of the Hop1-pT318 signal, the *pph3*Δ *pch2*Δ double mutant showed severe delay in meiosis progression (Figure 5F and G). The *pch2*Δ mutation suppresses pachytene arrest in the *zip1* mutant (San-Segundo and Roeder 1999; Borner *et al*. 2008). It, however, cannot suppress the delayed progression of meiosis in the *pph3*Δ mutant. Thus, this delay in *pph3*Δ can be caused by a mechanism different mechanism from that observed in the *zmm* mutant.

## Discussion

During meiotic recombination, the chromosome axis serves as an essential scaffold. Some components, including Mer2 and Rec8, are associated with pre-meiotic DNA replication. In contrast, the molecular mechanism underlying the formation of highly organized chromosomal axis structure, especially, the initial step of Hop1 recruitment onto meiotic chromatin, is not well understood. The results of this study support indications that PP4 may function in Hop1 recruitment onto chromatin in the initiation step of chromosome axis formation in early meiotic prophase-I.

### PP4 is involved in Hop1 recruitment onto chromatin independently from meiotic DSB-induced Tel1/Mec1 kinase activity

Significant delays in Hop1-Red1 loading were observed in *pph3* mutants, *pph3*Δ and *pph3-H112N*. In contrast, the timings of the production of Hop1 protein and the assembly meiotic cohesin component Rec8 in *pph3* mutants were indistinguishable from those in wild type. Delayed loading of Hop1 was also observed in the absence of Mec1 Tel1 activity and meiotic DSB formation. These results suggest PP4 is involved in Hop1 assembly before meiotic DSB formation. It has been reported that PP4 mutants show a delay in completing pre-meiotic DNA replication (Falk *et al*. 2010). Thus, our results cannot completely exclude the possibility that delayed completion of pre-meiotic DNA replication leads to subsequent delayed axis formation. It has been reported Rec8 loading is linked to pre-meiotic DNA replication (Kugou *et al*. 2009). There was a slight delay in Rec8 loading in the *pph3*Δ (but not in the *pph3-H112N* mutant), and this delay was less affected compared to Red1 loading (Figure 3B). From these results, we concluded that PP4 functions in promoting Hop1 loading during chromosome axis formation, an observation reached in light of the indirect pre-meiotic DNA replicative effects of its function as well.

During the formation of the meiotic SC structure, PP4 is also required for centromere-mediated chromosome pairing through Zip1 phosphorylation in a Mec1/Tel1-dependent manner (Falk *et al*. 2010). Both *pph3* mutants displayed significant delays in Hop1 assembly onto chromatin. However, this delay was Tel1/Mec1-independent (Figure 2A and B). Therefore, the observed Hop1 assembly delay would not be caused by compromised centromere pairing. In *pph3* mutants, the appearance of γH2A which is phosphorylated by Mec1/Tel1 kinase was observed at the same timing as that in wild type. By contrast, Tel1/Mec1-induced Hop1 phosphorylation (Hop1-pT318) was delayed. This result supports the idea that delayed Hop1 phosphorylation is caused by the delayed assembly of Hop1 onto the axes. Notably, *pph3* mutation-dependent delay in Hop1 phosphorylation was observed, not only in p*CLB2*-*MEC1*, but also in the *mec1*Δ backgrounds (Supplemental Figure 1A). This suggests that PP4 promotes timely recruitment of Hop1 onto chromatin independent of Mec1 (or Mec1 / Tel1) activity, rather than the indirect effects of Mec1-PP4-related premeiotic S-phase defects. Since Hop1 chromatin loading is one of the rate-determining factors that promotes meiotic DSB formation (Panizza *et al*. 2011), delayed DSB formation and Rad51 loading in the *pph3* mutant would also be caused by this delayed Hop1 loading onto the axes. In addition, our study has revealed the physical interaction between the Hop1 and a PP4 component, indicating that the Hop1 protein in the precipitates was not always phosphorylated (Figure 4A). These findings suggest that the Hop1-PP4 interaction may not only occur for enzymatic activity but may also regulate Hop1 assembly.

Another possibility is that delayed observation of Hop1 localization on the chromosome axes, in the absence of PP4, is unstable and thus rapidly removed from the axes. To assess this, we examined Hop1 localization in the absence of Pch2. Proper Hop1 assembly did not occur in 32% of the chromosomes in each nucleus of *pph3*Δ *pch2*Δ double mutant (Figure 5C). This strongly suggests that PP4 is involved in the recruitment phase of Hop1 and not in its stabilization on the chromosome axes. Conversely, 68% of chromosomes in the same nucleus displayed stable Hop1 assembly by *pch2*Δ, even in *pph3*Δ. These observations indicate that multiple pathways, beyond the PP4 alone, can promote Hop1 assembly.

### What molecular mechanism promotes PP4 induced Hop1 assembly?

The molecular mechanism by which PP4 promotes Hop1 assembly is still unknown. Furthermore, at this point, it is not clear whether the effect of PP4 on axis formation is a direct or indirect effect. Hop1 is a HORMA-domain protein. The dynamics of C-terminal “closure-motif” may be important for chromosome axis assembly and disassembly through Red1 interaction. In addition, substitution of K593 of Hop1 with alanine (*hop1-K593A*) results in severe defects in the Red1 interaction (West *et al*. 2018). We identified *hop1-S595L* mutant which showed hypomorphic Hop1 function defects such as normal spore viability and Hop1 assembly, but reduced Hop1 protein production, slightly delayed Hop1 phosphorylation at T318 and meiosis-I progression (Supplemental Figure 4A). We analyzed Hop1 assembly onto chromatin in the *pph3* within the *hop1-S595L* mutation background. The *pph3* and the *hop1-S595L* mutations showed a synergistic defect, thus *HOP1* and *PPH3* showed a genetic interaction in Hop1 assembly (Supplemental Figure 4B).

In addition, a delay in Hop1 assembly in the *pph3* mutant was also observed in the p*CLB2*-*MEC1 tel1*Δ and *spo11-Y135F* background, and Pph3 catalytic activity was required for timely recruitment of Hop1 onto chromatin (Figure 2A and B). These observations indicate PP4 dephosphorylation of unknown target protein before meiotic DSB formation is required for appropriately timed, Hop1 assembly onto chromatin (Figure 6). Since Hop1-PP4 physical interaction was detected simultaneously with Hop1 recruitment in the wild type (at 2 h; Figure 4A), Hop1 may be one of the most probable target for dephosphorylation by PP4 in the recruitment process. In contrast, the candidate protein kinase(s) regulating Hop1 assembly are DDK and/or CDK. However, the ATP analog sensitive *cdc7* allele (*cdc7-as3*) reportedly showed normal Hop1 production, despite a lack of Hop1 phosphorylation, in the presence of an ATP analog (Wan *et al*. 2006). The focus formation of Cdc28 (a component of CDK) on meiotic chromatin are dependent upon Hop1, Red1 and Rec8, but not on Mek1/Mre4. The analog sensitive allele *cdc28-as1* reportedly displayed compromised Rad51 focus formation and severe defects in Zip1 elongation. However it did not affect Red1 loading in the presence of an ATP analog (Zhu *et al*. 2010). These findings indicate that CDK and DDK functions are associated with Hop1 function. However, direct phosphorylation of Hop1 by CDK and/or DDK has not been reported. There are two predicted CDK phosphorylation motifs (S/T-P) in Hop1: S494 and S531 (Zhu *et al*. 2010). The presence of sequential phosphorylation by CDK followed by DDK in the sequence of serine/threonine residues in addition to the CDK phosphorylation motif has been described for related target proteins such as Mer2 (Wan *et al*. 2006) and Mps3 (RAO *et al*. 2020). Furthermore, the S494 of Hop1 overlaps this motif (T-S-^494^S-P). In addition, the possibility remains that Tel1/Mec1 kinase activity during the premeiotic S-phase is involved in the promotion of Hop1 loading.

**Figure 6.**
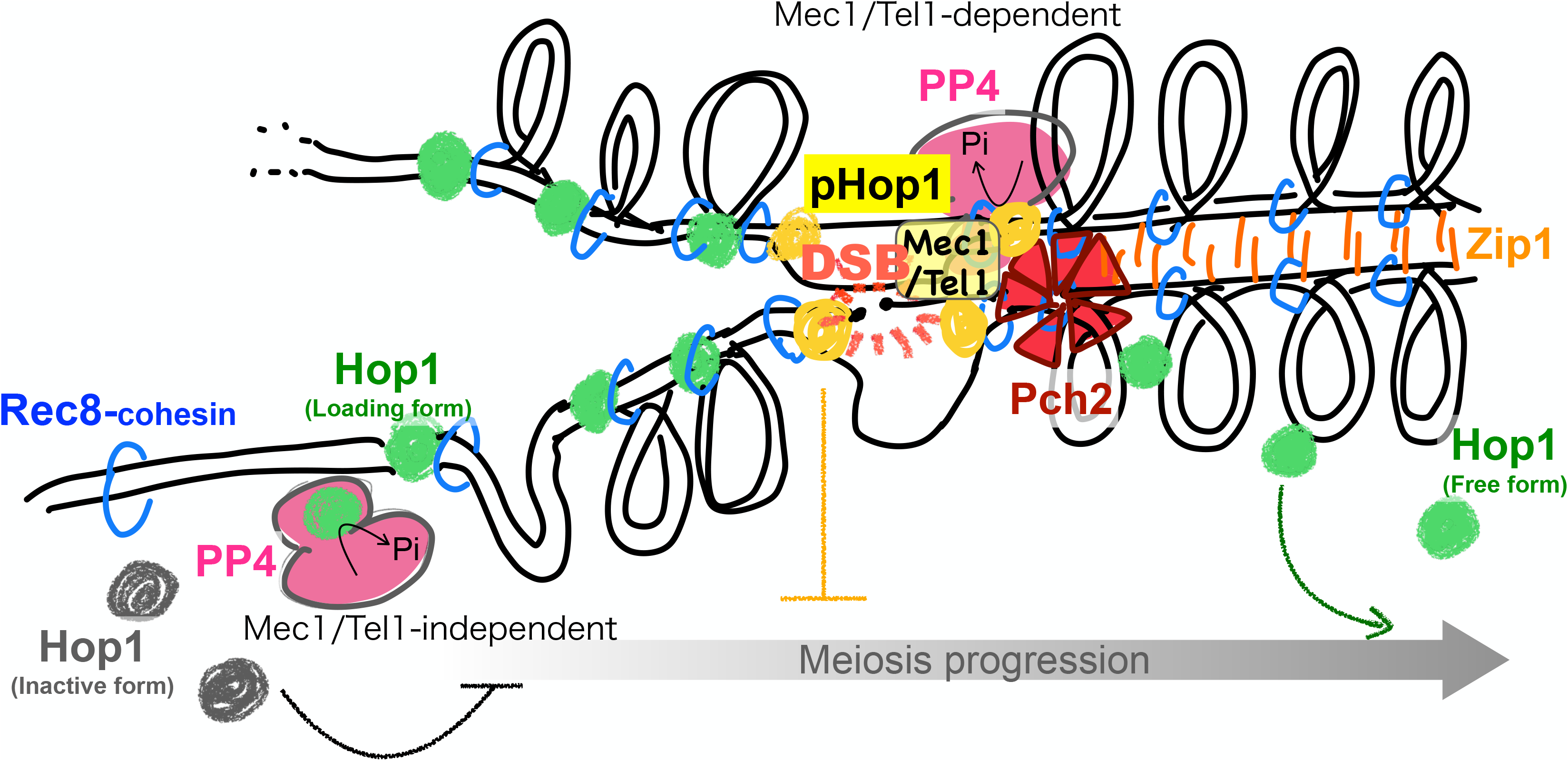
Model for PP4 involvement in Hop1 recruitment onto meiotic chromatin. PP4 plays multiple roles during meiotic prophase. First, independent of the introduction of meiotic DSBs and the function of Mec1/Tel1 activated by them, PP4 promotes Hop1 recruitment in the early phase of meiotic prophase. Until the onset of chromatin loading, Hop1 exists in an inactive form that may prevent miosis progression. After meiotic DSB formation, Hop1 is phosphorylated at the DSB site by Tel1/Mec1, and PP4 is involved in dephosphorylation of the phosphorylated Hop1. Meiotic recombination promotes homolog synapsis by exchanging Hop1 to Zip1 through the function of AAA+ Pch2 function.

### PP4 has an independent role from ZMM/SIC components in the prophase checkpoint

Pch2 activates the meiotic prophase checkpoint when meiotic DSB repair or chromosome synapsis is compromised in *zip1*, *zip2*, or *dmc1* mutant cells (San-Segundo and Roeder 1999; Borner *et al*. 2008). Prior studies have reported that the PP4 mutant shows a *zmm* mutant-like phenotype in several processes, including meiotic recombination, NCO/CO ratio aberration, temperature-sensitive meiosis progression, CO formation, and compromised CO interference (Borner *et al*. 2004; Falk *et al*. 2010). In addition, defective CO formation in *pph3*Δ is epistatic to *msh4*Δ (Falk *et al*. 2010). We also observed compromised CO and elevated NCO formation via HD analysis (Figure 1C). These are commonly observed in *zmm* mutants (Shinohara *et al*. 2008). Moreover, although *pch2*Δ suppresses meiotic prophase arrest of *zmm* mutants (San-Segundo and Roeder 1999), it could not suppress the delayed meiosis progression of the *pph3*Δ mutant. These findings suggest that delayed meiosis progression is caused by a different mechanism from that of *zmm* or *dmc1* mutants. It was rrecently eported that Pch2, in collaboration with Hop1, mediates the meiotic prophase checkpoint (Raina and Vader 2020; Herruzo *et al*. 2021). In *pph3* mutants, compromised synapsis as well as inefficient meiotic recombination easily activate the meiotic prophase checkpoint because of enhanced Mec1/Tel1 signaling.

However, the checkpoint signal was not restored even in the absence of Pch2. In addition, we also observed delayed meiosis progression in *pph3* mutants without meiotic DSB formation in the *spo11-Y135F* background (Supplemental Figure 2A). This could be caused by insufficient Hop1 loading on the chromosome axis or an accumulation of unbound inactive form of Hop1 protein. Further, PP4 might be involved in inactive to active conversion of Hop1 before chromatin loading, thus ensuring proper axis construction (Figure 6).

## Materials and Methods

### Yeast strains and media

All the yeast strains and their genotypes are listed in Supplemental table 1. We used isogenic *S. cerevisiae* SK1 background NKY1551 derivatives (Storlazzi *et al*. 1996). *PPH3-13MYC* was constructed using a polymerase chain reaction–based tagging method (Bahler *et al*. 1998). *pph3-H112N* was constructed vis two-step gene replacement. YPAD (1% yeast extract, 2% Bacto peptone, 0.01% adenine, 2% glucose), SPM (0.3% potassium acetate, 0.03% raffinose), and DISS plates (YPAD medium containing 2.4% Bacto Agar) were used.

### Yeast meiosis time course

The experiments were performed as described previously (Shinohara *et al*. 2003). Meiosis progression was analyzed by counting the number of nuclei in each ascus at each time point. Nuclei were visualized by staining with 4’, 6-diamidino-2-phenylindole, dihydrochloride (DAPI) and observed under an epifluorescence microscope (AxioSkop2, Carl Zeiss). More than 100 nuclei were analyzed at each time point.

### Cytological analysis

Immunostaining of yeast meiotic nuclear spreads was performed as previously described (Shinohara *et al*. 2015). Stained samples were observed under epifluorescence using the Axioskop 2 microscope equipped with light-emitting diode fluorescence light sources (X-Cite; Excelitas Technologies) and a 100× objective (AxioPlan, NA1.4, Carl Zeiss). Images were captured with a CCD camera (Retiga; Qimaging) and processed using iVision (BioVision Technologies) and Photoshop (Adobe). The antibodies used in these assays were anti-Hop1 (Iwasaki *et al*. 2016) (guinea pig, 1:500), anti-Hop1-pT318(Iwasaki *et al*. 2016) (rabbit, 1:500), anti-Red1 (chicken, 1:500) (Shinohara *et al*. 2008) and anti-Rec8 (rabbit, 1:1000) (Zhu *et al*. 2010). Nuclei containing more than five foci were counted as focus-positive.

### Southern blot to detect meiotic DSBs and recombination intermediates

Genomic DNAs prepared from yeast meiosis time course experiments were digested with *Mlu*I, *Bam*HI, and *Xho*I for heteroduplexes (HDs), and *Pst*I for meiotic DSBs. Probes for Southern blotting included: Probe 155 for HD and Probe 291 for DSB detection (Storlazzi *et al*. 1996; Shinohara and Shinohara 2013). They were labeled with α-^32^P-dATP (NEG512H, PerkinElmer) using the random primer method (Random Primer 6, NEB S1230S, Klenow fragment (3’-5’ exo-, M0210) and visualized by using FLA7000 (Cytiva). The visualized signals were analyzed using Image Quant TL software (Cytiva) and images were processed using Photoshop (Adobe).

### Western blotting

Western blotting was performed as described previously (Shinohara *et al*. 2015). Meiotic yeast cell extracts were prepared by cell disruption using a Multi-Beads Shocker (YASUIKIKAI) after trichloroacetic acid treatment. The primary antibody signal was visualized using Alexa Fluor 680–labeled secondary antibodies (Thermo Fisher Scientific) or IRDye 800CW–labeled secondary antibodies (LI-COR Biosciences) using an Odyssey infrared imaging system (LI-COR Biosciences). Images were processed using Photoshop (Adobe). The primary antibodies used in these assays were anti-γH2A (anti-γH2A, AF2288, R&D Systems, 1:1000), anti-Myc (MC045, Nacalai Tesque, 1/1000), anti-Hop1 (guinea pig, 1/5000)(Iwasaki *et al*. 2016), anti-Hop1-pT318 (rabbit, 1/5000) (Iwasaki *et al*. 2016), anti-Rec8 (rabbit, 1/10000) (Zhu *et al*. 2010), anti-Red1 (chicken, 1/3000) (Shinohara *et al*. 2008) and anti-α-tubulin (YL1/2, Santa Cruz Biotechnology, 1/3000). Tubulin signals were visualized using the LI-COR Bioscience system or alkaline phosphatase stain (BCIP-NBT solution kit, Nacalai Tesque) on the same membrane on which other meiotic proteins were detected.

### Immunoprecipitation

The immunoprecipitation assay was performed as previously described with minor modifications (Shinohara *et al*. 2008). Psy2-13Myc was immunoprecipitated using anti-Myc antibody (MC045, Nacalai Tesque) These were bound to Dynabeads Protein G (Thermo Fisher Scientific) from meiotic yeast cell lysate prepared using a Multi-Beads Shocker (YASUIKIKAI) in lysis buffer (50 mM HEPES-NaOH, pH7.5, 150 mM NaCl, 20% glycerol, 1 mM Sodium orthovanadate, 60 mM β-glycerophosphate, 0.1% Nonidet P-40, 1:100 dilution of protease inhibitor cocktail (SIGMA-Aldrich) at the indicated time points in Figure 4A and 4B. Psy2-13Myc, Hop1, Red1 and Rec8 proteins in the precipitates and WCEs were detected as described in the western blot section.

### Tetrad analysis

Tetrad analysis was performed as described previously (Challa *et al*. 2019). Parental haploid strains were mated for 12 h on YPAD plates at 30°C and then transferred onto SPM plates. After incubation at 30°C for 48 h, tetrads were dissected onto DISS plates and incubated for 48 h.

## Data availability

Strains and plasmids are available upon request.

The authors affirm that all data necessary for confirming the conclusions of the article are present within the article, figures, and tables.

## Acknowledgements

We are grateful to Dr. K. Matsuzaki, Dr. F. Klein, Dr. S. Gasser, and the members of the Shinohara lab for their helpful discussions. We thank Ms. A. Murakami and Ms. A. Kitada for their excellent technical assistance. K.L. was supported by a Fellowship from the Institute for Protein Research, Osaka University. We would like to especially thank Dr. A. Shinohara and Dr. K. Matsuzaki for critical reading of the manuscript. This work was supported by JSPS KAKENHI (#15H05973, #19K22402) and Takeda Science Foundation and was performed in part under the Cooperative Research Program of the Institute for Protein Research, Osaka University, CR-17-05.

---

**Supplemental Figure 1.**
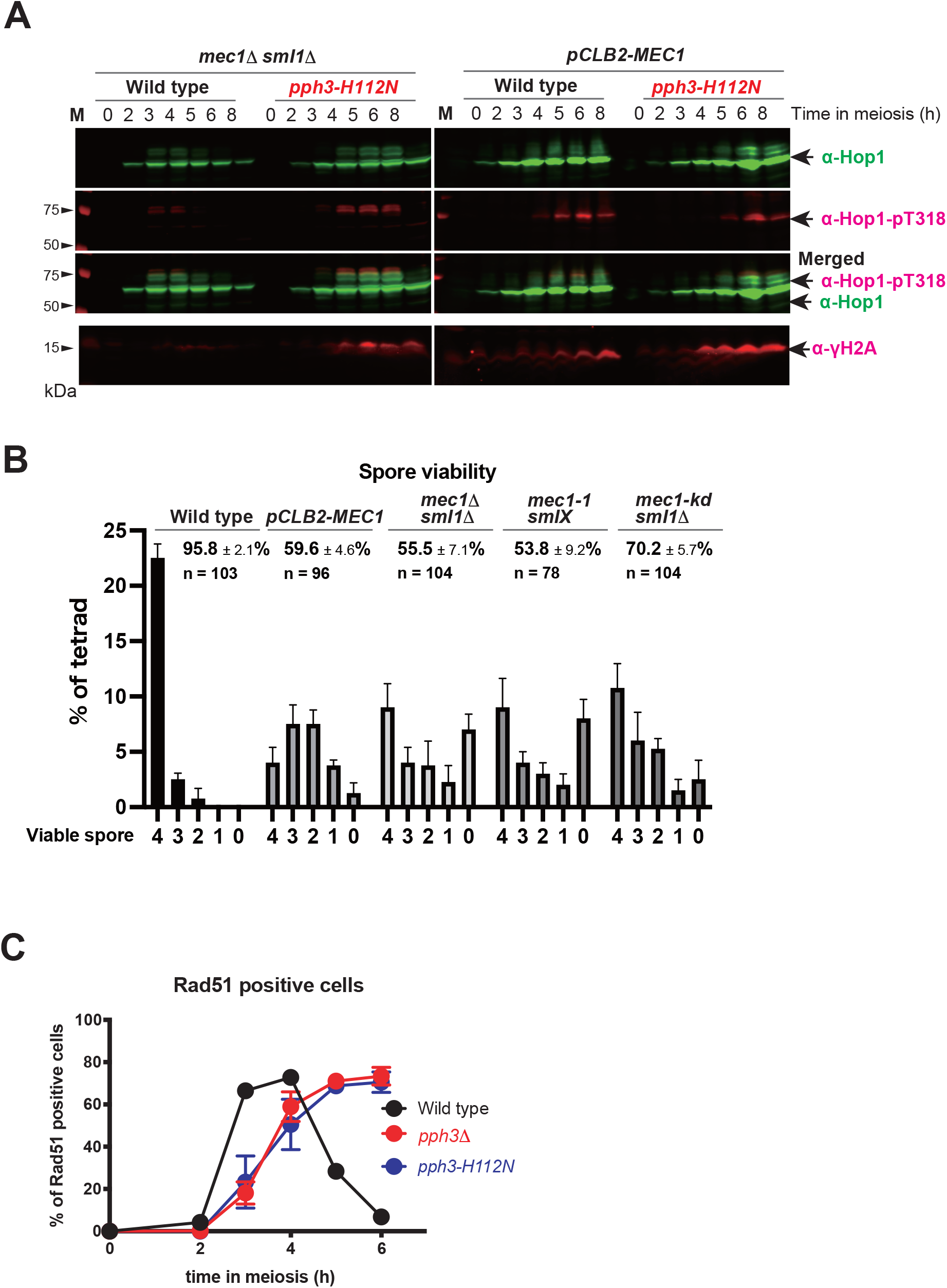
Supplemental data for Figure 1. (A) Representative western blot images of Hop1 (green, top), phosphorylation of Hop1 at T318 (Hop1-pT318, red, middle), merged images of Hop1 and Hop1-pT318, and phosphorylation of histone H2A (γH2A, red, bottom) at the indicated time point during meiosis in *mec1*Δ *sml1*Δ (MSY3643/3644), *mec1*Δ *sml1*Δ *pph3-H112N* (MSY6560/6565), *pCLB2-MEC1* (MSY6551/6552), and *pCLB2-MEC1 pph3-H112N* (MSY6574/6576). M denotes molecular weight markers. (B) Spore viability and the distribution of 4-, 3-, 2-, 1-, and 0-viable spores in wild type (MSY833/831), *pCLB2-MEC1* (MSY6551/6552), *mec1*Δ *sml1*Δ (MSY3643/3644), *mec1*-1 *sml1X* (MSY398/399), and *mec1*-*kd sml1*Δ (MSY4413/4415). n: number of asci analyzed. (C) The graph of the kinetics of Rad51 focus assembly on meiotic nuclear spreads in wild type (black, NKY1303/1543), *pph3*Δ (red, MSY5632/MSY5634), and *pph3-H112N* (blue, MSY6219/6220). More than 100 cells were analyzed at each time point. Error bars show the mean ± standard deviation of more than three trials.

**Supplemental Figure 2.**
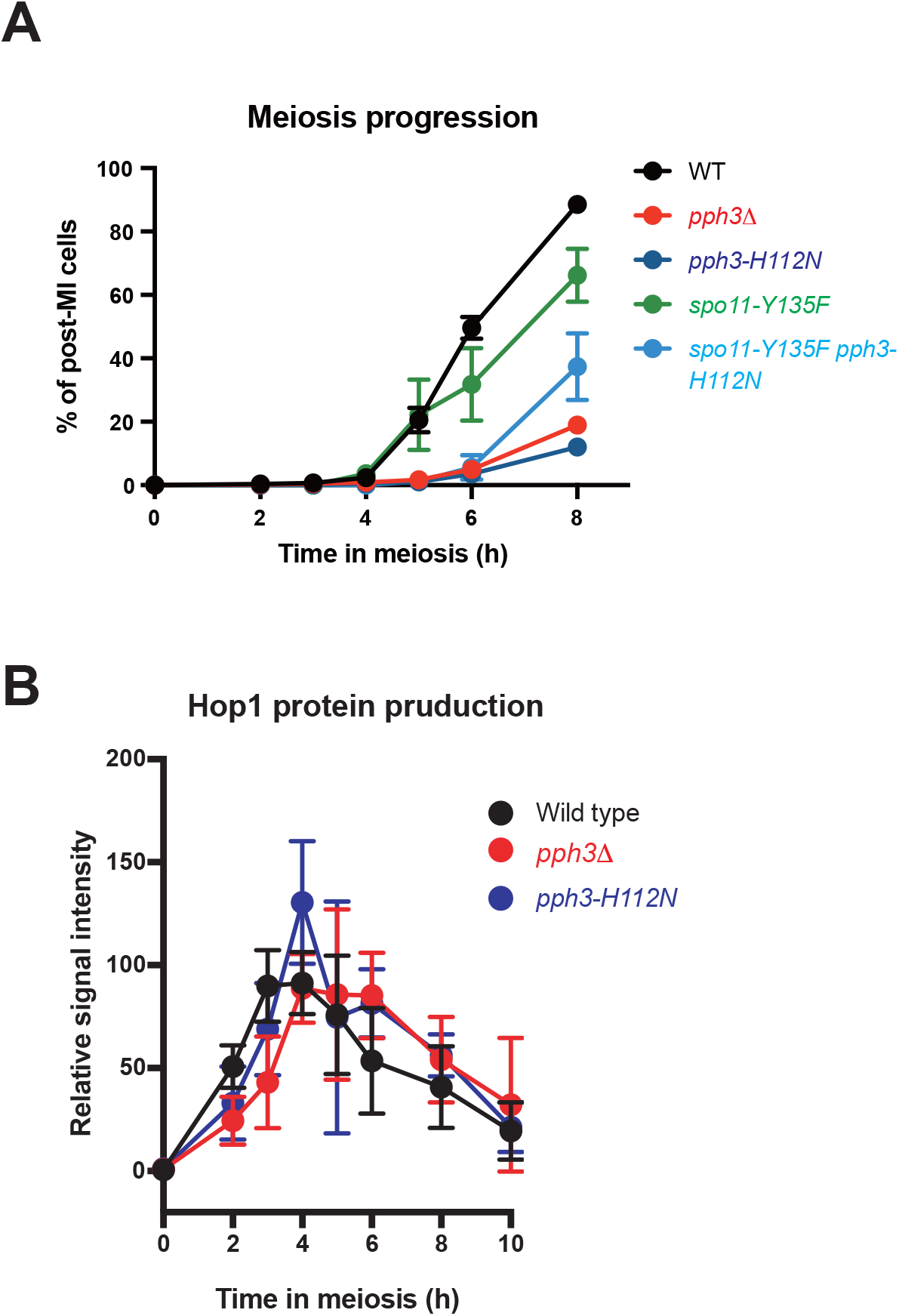
Supplemental data for Figure 2. (A) Meiosis progression in wild type (NKY1303/1543), *pph3Δ* (MSY5632/5634), *pph3-H112N* (MSY6219/6220), *spo11-Y135F*(MSY4139/4141), and *spo11-Y135F pph3-H112N* (MSY6568/6570) mutants. Frequencies of nuclei with more than two DAPI-stained bodies were plotted. More than 200 cells were analyzed at each time point. (B) Graphical depiction of the average number of relative Hop1 protein signals from western blot at each time point. The numerical values are shown as a percentage of the peak amount of signal in the wild type. Error bars show the mean and ± standard deviation of two independent trials.

**Supplemental Figure 3.**
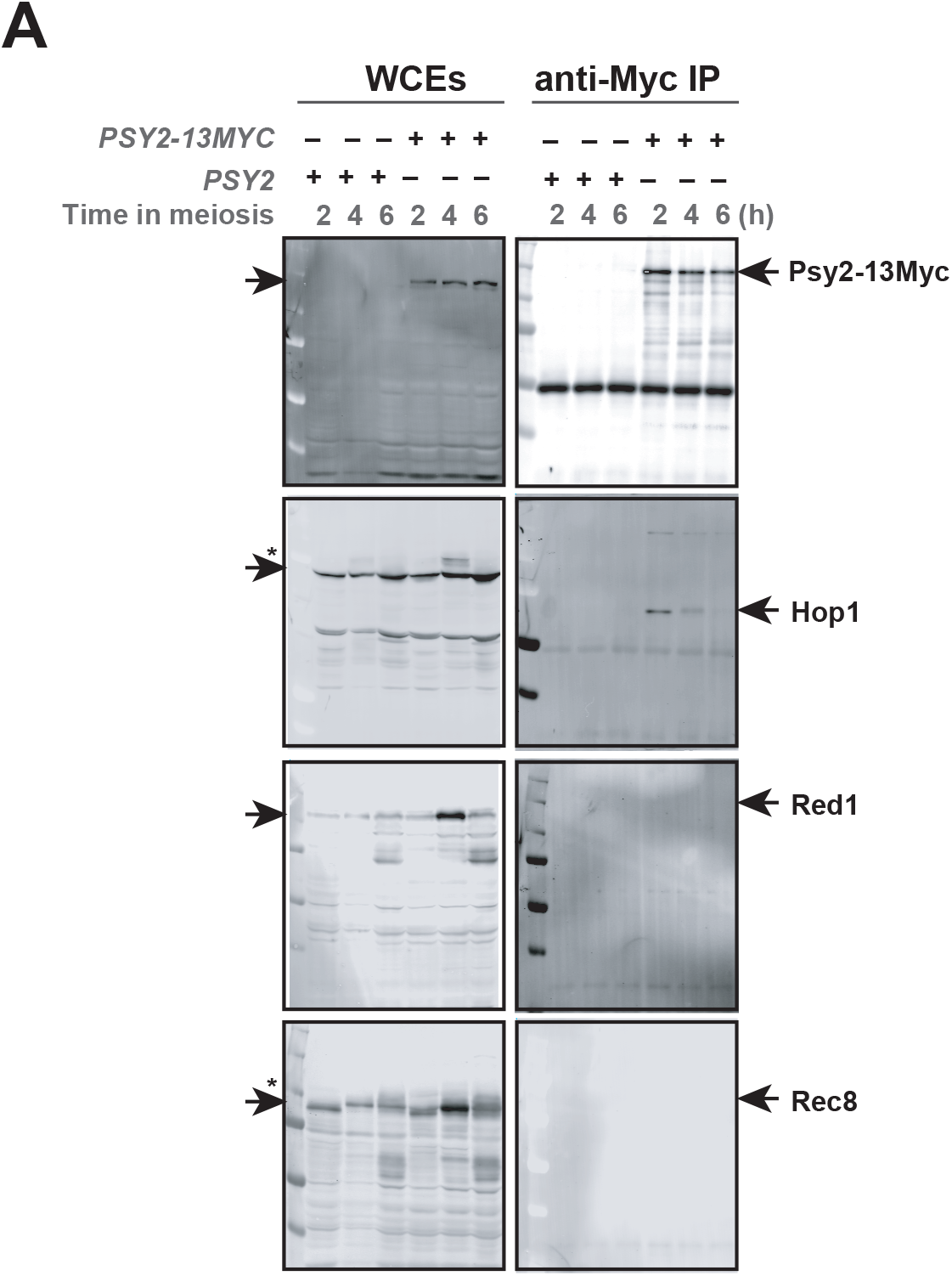
Supplemental data presentation for Figure 4. (A) Whole membrane images of western blot of immunoprecipitates (IP) and whole cell extracts (WCEs) exhibited in Figure 4A.

**Supplemental Figure 4.**
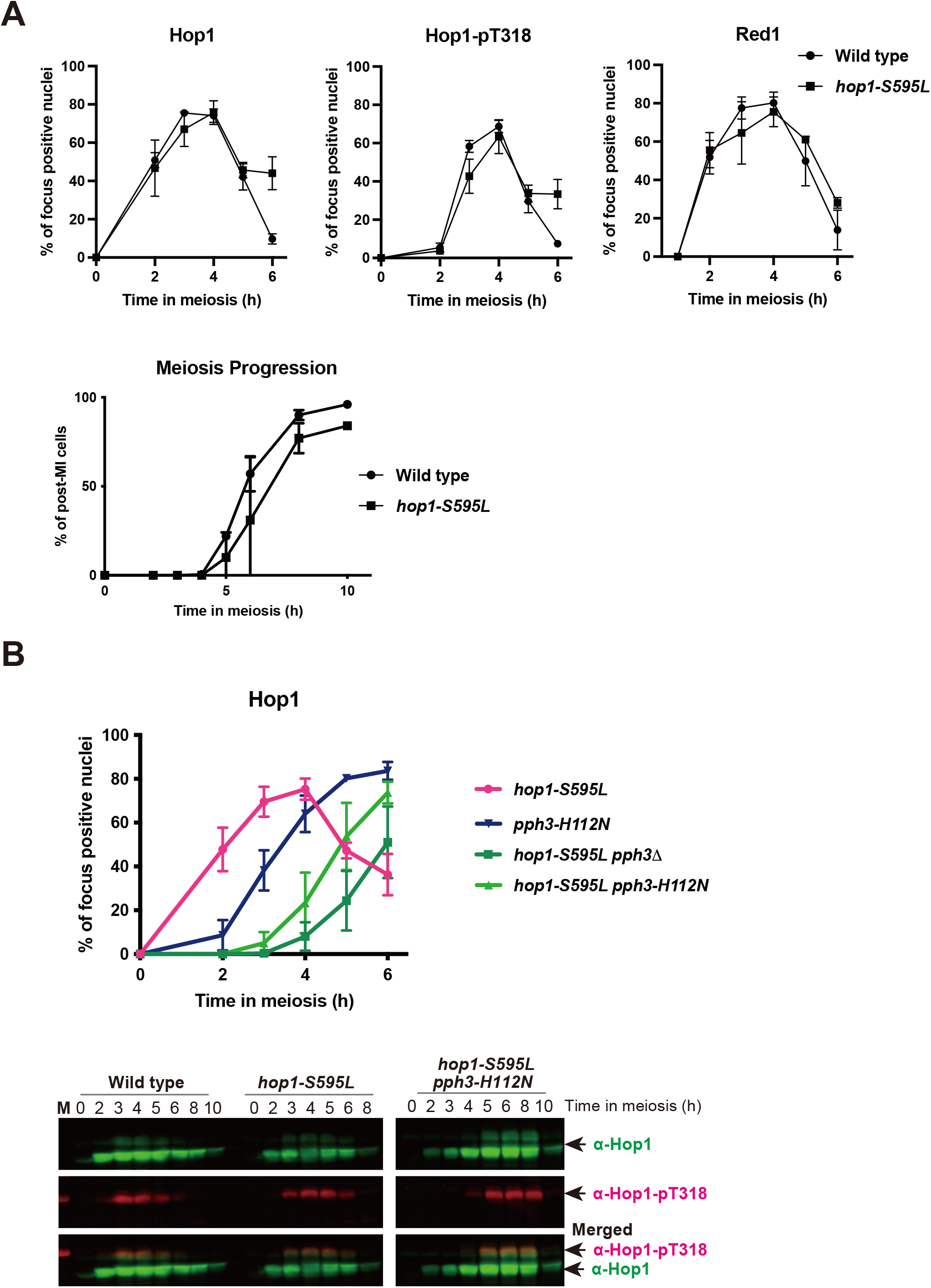
Characterization of *hop1-T595L* as a hypomorph allele of *HOP1*. (A) Graphical depiction of the kinetics of Hop1, Hop1-pT318, and Red1 focus assembly on meiotic nuclear spreads in the indicated strains, wild type (MSY833/831) and *hop1-T595L* (MSY6314/6319) (upper). More than 100 cells were analyzed at each time point. Error bars show the mean ± standard deviation of more than three trials. Meiosis progression in wild type (MSY833/831) and *hop1-T595L* (MSY6314/6319). Frequencies of nuclei with more than two DAPI-stained bodies were plotted (lower). More than 200 cells were analyzed at each time point. (B) Graphical depiction of the kinetics of Hop1 focus assembly on meiotic nuclear spreads in the indicated strains, *hop1-T595L* (pink, MSY6314/6319), *pph3*Δ *hop1-T595L* (red, MSY6544/6546), and *pph3-H112N hop1-T595L* (green, MSY6541/6543). Values of *pph3-H112N* (dark blue) are from the same data shown in Figure 2A. More than 100 cells were analyzed at each time point. Error bars show the mean ± standard deviation of more than three trials. Representative western blot images of Hop1 (green, top), phosphorylation of Hop1 at T318 (Hop1-pT318, red, middle), and merged images of Hop1 and Hop1-pT318 at the indicated time point during meiosis in wild type (MSY833/831), *hop1-T595L* (MSY6314/6319), *pCLB2-MEC1* (MSY6551/6552), and *pCLB2-MEC1 pph3-H112N* (MSY6574/6576). M denotes molecular weight markers.

**Supplemental Table-1.**
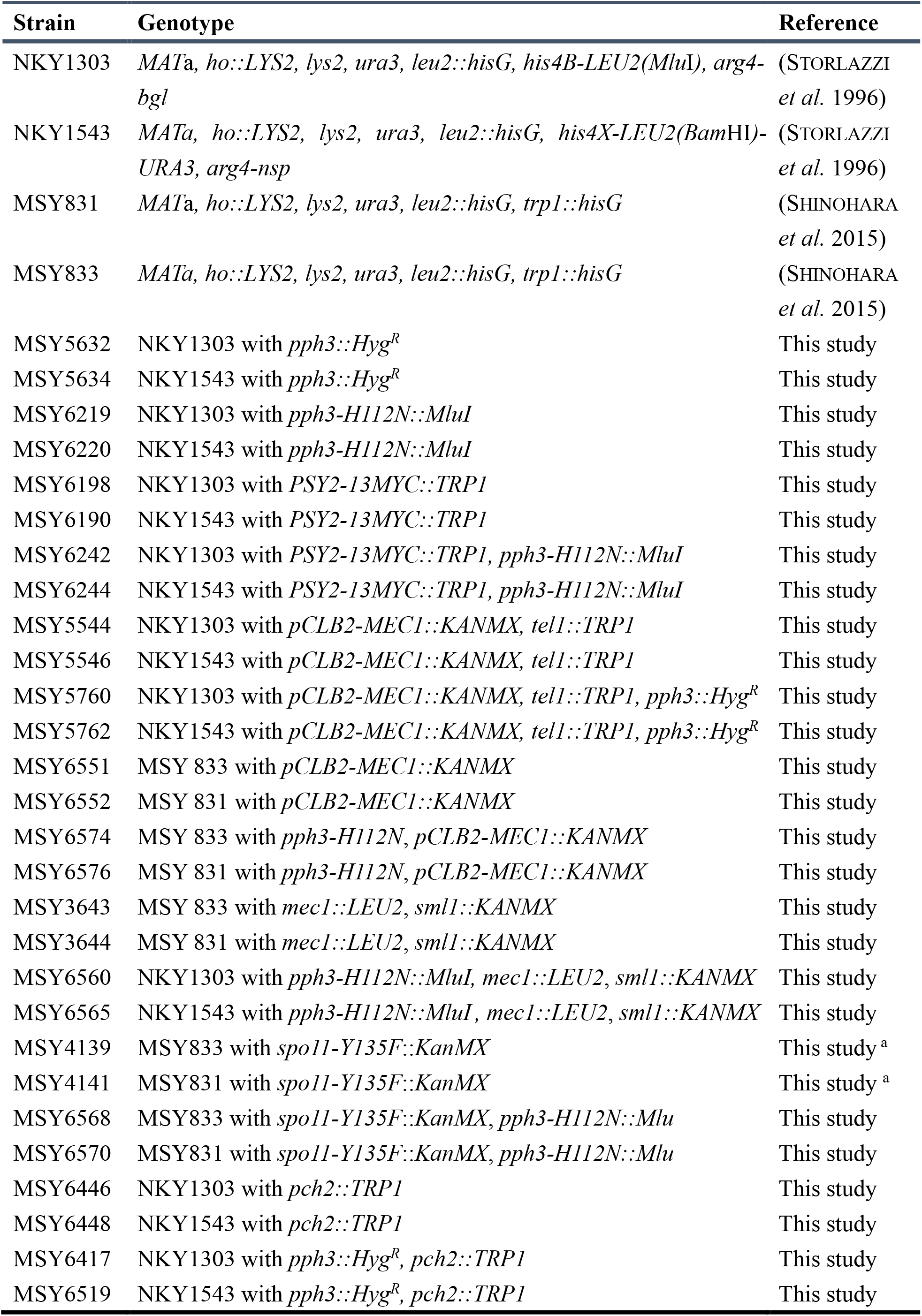
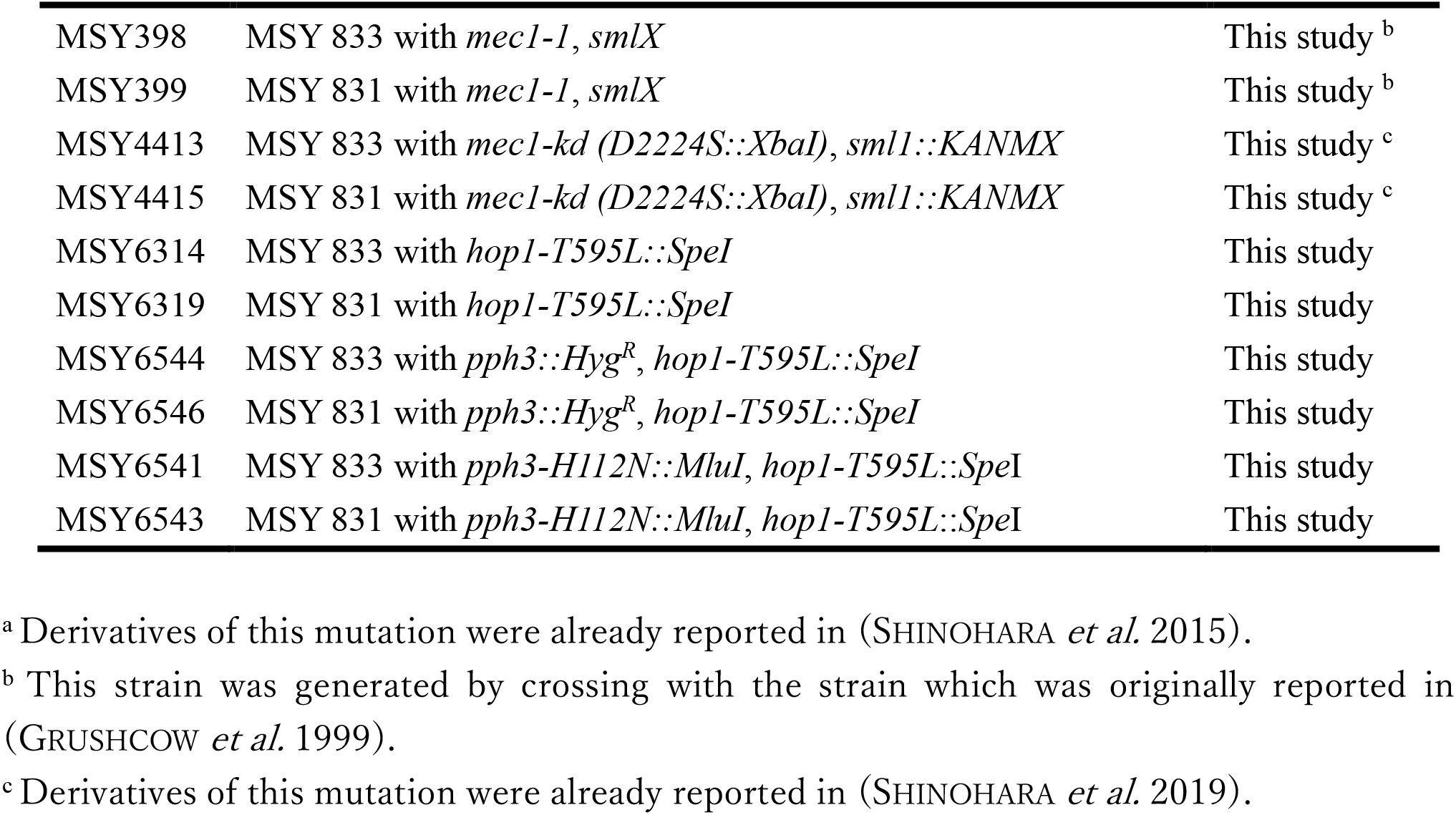
Strain list

## Notes

### Competing Interest Statement

The authors have declared no competing interest.

### Summary of Updates

In this version, we have added analyses for characteriing the pCLB2-MEC1 allele to compare the meiotic phenotype with that of other MEC1 defective alleles, such as mec1Δ (sml1Δ), mec1-1 (smlx), and mec1-kd (sml1Δ). We have also added analyses for characterization of the newly identified hop1 hypomorph allele, hop1-T595L, and data of Hop1 assembly in hop1-T595L pph3 double mutants to show genetic interaction between HOP1 and PPH3 in Hop1 assembly. In addition, we have added analyses for Hop1 assembly in spo11-Y135F, a catalytic-dead allele. This helped us to clarify whether Tel1/Mec1 function, activated by meiotic DSB formation, is dispensable from PP4 function in Hop1 loading. In contrast, we found that we could not improve upon the fact that there is no residual activity of Mec1 in pCLB2-MEC1. Finally, we removed Tel1/Mec1-independent from the title as suggested.

